## Supplementary Information for "A Corporative Language Model for Protein–Protein Interaction, Binding Affinity, and Interface Contact Prediction"

#### Supplementary Texts

##### Text S1. Inter-protein attention matrix analysis of PPLM and ESM2

To further investigate what is captured by PPLM beyond existing single-protein language models, we conducted a representation-level comparison between PPLM and ESM2. Specifically, we examined how their inter-protein attention matrices relate to true inter-chain contacts under two complementary settings: (i) **Unsupervised**, where the last-layer inter-protein attention was averaged across heads and layers to directly predict contact likelihoods; and (ii) **Linear-probe**, where a single frozen linear layer was trained on all-layer inter-protein attentions using the training set of PPLM-Contact (5 epochs, learning rate = 0.001).

Performance was evaluated on the Homodimer300 and Heterodimer99 test sets in terms of Top-*k* inter-protein contact precision ($k \in \{1, 5, 10, 25, 50, 100, L/10, L/5, L/2, L\}$, where $L$ is the length of the shorter protein in each complex). Paired Wilcoxon signed-rank tests (one-sided, PPLM > ESM2) were used to assess statistical significance. Error bars in **Figure S2** represent the standard error (SE) across complexes, and significance levels are annotated as $p < 0.05$ (*), p < 0.01 (**), and $p < 0.001$ (***).

On the Homodimer300 test set, PPLM shows consistent and statistically significant gains over ESM2 across all Top-*k* thresholds. In the unsupervised last-attention setting, the mean improvements (PPLM−ESM2, percentage points) are +7.33 (Top-1), +6.87 (Top-5), +5.23 (Top-10), +4.23 (Top-25), +3.18 (Top-50), +2.03 (Top-100), +4.53 (Top-*L*/10), +3.59 (Top-*L*/5), +2.52 (Top-*L*/2), and +1.78 (Top-*L*); all are significant according to paired Wilcoxon tests (***, $p < 0.001$). With the single linear probe, gains remain robust—+6.67, +6.40, +5.57, +4.47, +3.31, +2.20, +4.43, +4.14, +2.37, +1.49 for Top-1 to Top-*L*—with statistical significance at at $p < 0.01$ (**) or $p < 0.001$ (***). Specifically, PPLM achieves 6.45% and 11.68% Top-*L* precision in the unsupervised and linear-probe settings, representing 38.4% and 14.7% relative improvements over ESM2 (4.66% and 10.18%). At the per-complex level (Top-*L* evaluation), PPLM outperformed ESM2 on 195 homodimers, tied on 24, and underperformed on 81 in the last-layer-attention setting; the corresponding numbers for the linear-probe setting are 161, 20, and 119.

On the Heterodimer99 test set, in the unsupervised last-attention setting, PPLM performs lower than ESM2 at Top-1 (−1.52 points) but surpasses it from Top-10 onward (e.g., +1.77 at Top-10, +1.03 at Top-*L*/5, +0.41 at Top-*L*). Correspondingly, Wilcoxon tests are not significant for Top-1, Top-5, and Top-*L*/10, but become significant from Top-25 onward (except Top-*L*/10), indicating a gradual emergence of representational advantage as k increases. With the single linear probe, PPLM improves by +4.04 (Top-1), +2.22 (Top-5), +2.22 (Top-10), +2.42 (Top-25), +1.76 (Top-50), +1.67 (Top-100), +2.32 (Top-*L*/10), +1.95 (Top-*L*/5), +1.89 (Top-*L*/2), and +1.74 (Top-*L*). Statistical significance is observed at Top-25, Top-50, Top-100, Top-L/10, Top-L/2, and Top-*L* ($p$ < 0.05 / 0.01 / 0.001, indicated as */**/***), while Top-1/5/10 and Top-*L*/5 show no formal significance but still exhibit notable average precision gains. Specifically, PPLM achieves 2.76%

and 6.44% Top-$L$ precision in the unsupervised and linear-probe settings, representing 17.5% and 27.0% relative improvements over ESM2 (2.35% and 4.70%). At the per-complex level, PPLM outperformed ESM2 on 54 heterodimers, tied on 23, and underperformed on 22 in the last-layer-attention setting; for the linear-probe setting, the corresponding numbers are 51, 28, and 20.

These results demonstrate that the improvement provided by PPLM is broadly distributed across both homodimer and heterodimer proteins, with consistent advantages in the majority of cases. Although the gains are smaller and less significant at Top-1 for heterodimers in the unsupervised setting, PPLM consistently achieves higher precision as $k$ increases. These findings indicate that PPLM's inter-protein attention encodes more explicit and linearly decodable patterns of inter-protein interactions than ESM2, confirming that paired-protein language modeling captures deeper relational representations beyond single-chain embeddings.

#### Text S2. Ablation analysis of PPLM-PPI

To assess the influence of network-level pooling operations and input feature compositions on cross-species PPI prediction, we performed two groups of ablation experiments for PPLM-PPI. Each variant was evaluated on the *M. musculus*, *D. melanogaster*, *C. elegans*, *S. cerevisiae*, and *E. coli* test sets, and we report the mean accuracy ± 95 % confidence intervals (CI) across three random seeds.

Seven pooling configurations were compared, combining mean, max, and min operations in different ways. As shown in **Figure S4A**, the mean–max pooling consistently achieved the best or near-best AUPRC across all species, reaching 0.921 ± 0.002 in *M. musculus*, 0.906 ± 0.002 in *D. melanogaster*, and 0.884 ± 0.002 in *C. elegans*. Removing either the mean or max component led to noticeable declines (e.g., mean only: 0.914 ± 0.002; max only: 0.903 ± 0.002 in *M. musculus*). Among single pooling operations, mean pooling achieved the highest accuracy, while min pooling performed the worst across species, likely due to its sensitivity to weak or noisy signals. These trends are consistent across all five species, confirming that combining mean and max pooling provides a balanced representation of global and salient pairwise features. The narrow confidence intervals (< 0.01) demonstrate stable performance across random seeds.

Under the optimal single mean-pooling configuration, we further examined the contributions of different feature types, including PPLM-derived inter-attention, intra-attention, and embedding features. As shown in **Figure S4B**, the full model integrating all three components achieved the highest AUPRC across all species (e.g., 0.914 ± 0.002 in *M. musculus*, 0.886 ± 0.005 in *D. melanogaster*, and 0.853 ± 0.003 in *C. elegans*). Removing any attention stream markedly reduced performance—for instance, the inter-attention + intra-attention variant dropped to 0.560 ± 0.001 in *M. musculus* and 0.586 ± 0.007 in *D. melanogaster*—indicating that attention signals alone without embeddings are insufficient. Models retaining embeddings (e.g., *intra-attention + embedding* or *embedding only*) maintained substantially higher accuracy than those without, demonstrating that embeddings contribute most strongly to predictive performance. Models using a single attention type (either intra- or inter-attention alone) produced the lowest accuracies (~0.52–0.69).

Across both analyses, all confidence intervals remained below 1%, supporting the statistical robustness of the observed trends. Collectively, these experiments demonstrate that the mean–max pooling strategy and the combined inter-attention + intra-attention + embedding feature set are critical for achieving robust and generalizable cross-species PPI prediction. Detailed numerical results for all metrics are listed in **Table S2-3**.

**Text S3. Inductive generalization analysis under sequence-similarity stratification for PPI prediction**

To evaluate whether the performance of PPLM-PPI could be affected by shortcut learning and to directly assess its inductive generalization ability, we conducted a sequence-similarity–stratified analysis across all five non-human test species (mouse, fly, worm, yeast, and E. coli). For every protein in each test organism, we computed its maximum sequence identity to any human protein in the training set (alignment coverage $\geqslant$ 0.8). Test interaction pairs were then grouped into seven identity bins: (0, 0.5], (0.5, 0.6], (0.6, 0.7], (0.7, 0.8], (0.8, 0.9], (0.9, 1.0), and [1.0], according to the higher identity of its two constituent proteins, which reflects the maximal similarity between any protein in the pair and the training set. For each bin, we evaluated the AUPRC of PPLM-PPI and all baseline models (TUN-A, ESM-based PPI, D-SCRIPT, and Topsy-Turvy), and recorded the negative-to-positive sample ratio to quantify class imbalance (**Supplementary TableS6 and Supplementary Figure S5**).

Across species, the lowest-identity (0, 0.5] bin typically contains the strongest class imbalance, with negative–positive ratios ranging from 8.5:1 (E. coli) to 86.4:1 (mouse). Higher-identity bins are progressively less imbalanced, except for the small [1.0] bin in D. melanogaster, which is highly skewed due to limited sample size. This pattern indicates that raw AUPRC in extremely low-identity bins reflects the combined effect of homology and class imbalance rather than shortcut learning.

Despite large differences in class imbalance and sequence identity, PPLM-PPI maintains robust performance across all bins. In all five species, PPLM-PPI achieves high AUPRC for identity > 0.5 and does not exhibit degradation when identity decreases. In D. melanogaster and C. elegans, PPLM-PPI even attains higher AUPRC values in the (0.6, 0.9] bins than in the (0.9, 1.0) bin, showing that performance does not strictly track sequence similarity to the human training set. In the E. coli dataset, where all test proteins exhibit sequence identities below 0.8 relative to the human training set, PPLM-PPI also continues to deliver strong predictive accuracy.

Crucially, PPLM-PPI consistently outperforms all baseline methods in nearly every identity bin, including the most remote (0, 0.5] range. In this bin, PPLM-PPI improves AUPRC over TUN-A by 26.0%, 12.7%, 13.9%, 33.4%, and 13.3% on the mouse, fly, worm, yeast, and E. coli datasets, respectively. Similar or larger improvements are observed relative to ESM-based PPI, D-SCRIPT, and Topsy-Turvy. These results demonstrate that the advantage of PPLM-PPI is greatest in the most challenging remote-homology regime, where shortcut learning—if present—would be least effective.

Taken together, this similarity-stratified evaluation shows that (i) the dependence of AUPRC on identity is largely driven by underlying class imbalance and biological conservation and is shared by all models; (ii) PPLM-PPI maintains strong and stable predictive accuracy across sequence-identity ranges; and (iii) the method does not rely on protein-identity shortcuts, but instead generalizes robustly to unseen proteins in a strict cross-species inductive setting.

#### Text S4. Statistical analysis of PPLM-Contact compared with baseline methods

To quantitatively assess the performance improvements achieved by PPLM-Contact over existing inter-protein contact prediction methods, we conducted a comprehensive statistical analysis across four benchmark datasets (Homodimer300, Heterodimer99, CASP_Homo43, and CASP_Hetero20). For each target, we computed the Top-L inter-protein contact precision for PPLM-Contact and for each baseline model, enabling paired comparisons at the per-target level. Statistical significance was evaluated using paired Wilcoxon signed-rank tests, and effect sizes were quantified using paired Cohen's *d*. To correct for multiple hypothesis testing, *p*-values across all applicable datasets were adjusted using the Benjamini–Hochberg false-discovery-rate (FDR) procedure ($\alpha$ = 0.05). All statistical results are reported in **Supplementary Tables S11–S12**.

***Using distance maps derived from experimental monomer structures***, PPLM-Contact exhibits consistent and substantial improvements over every baseline across all four datasets. Against PLMGraph-Inter, PPLM-Contact achieves large effect sizes (Cohen's d = 0.57–1.04) and highly significant improvements ($q$ = $4.23\times10^{-38}$ to $9.56\times10^{-3}$) on all datasets, including the challenging CASP sets. Compared with DeepInter, a strong learning-based baseline, PPLM-Contact achieves medium effect sizes on the two large benchmarks (Cohen's d ≈ 0.43) and remains superior across all datasets. The only non-significant case occurs on CASP_Homo43, where the modest sample size yields a small effect (Cohen's d = 0.15, $q$ = 0.074); however, PPLM-Contact still attains higher mean precision, indicating a genuine performance advantage despite limited statistical power. Against CDPred, a representative structure-driven model, PPLM-Contact shows substantial gains in every dataset, with large effect sizes (Cohen's d = 0.57–0.87) and all FDR-corrected $q$-values below 0.01. For DeepHomo2.0—specialized for homodimers—PPLM-Contact demonstrates very large effects on both homodimer datasets (e.g., Homodimer300: $d$ = 1.08, $q$ = $6.22\times10^{-43}$). Finally, the strongest improvements are observed against GLINTER, with extremely large effect sizes (Cohen's d = 0.96–1.59) and extraordinarily small FDR-adjusted $q$-values (as low as $5.18\times10^{-47}$), demonstrating decisive gains across both homo- and heterodimer test sets.

***Using distance maps derived from AlphaFold2-predicted monomer structures***, PPLM-Contact achieves statistically significant improvements over all baseline methods across all four datasets. The effect sizes range from medium to very large (Cohen's $d$ = 0.35–1.25), and every comparison attains significance after FDR correction ($q < 0.05$). Notably, PPLM-Contact maintains strong gains over PLMGraph-Inter and CDPred, shows consistent improvements over DeepInter even on the smaller CASP sets, and continues to outperform DeepHomo2.0 with very large effects on homodimers (e.g., $d \approx 0.84$–0.89). The largest margins are observed against GLINTER, with effect sizes exceeding 1.0 and $q$-values as low as $2.26\times10^{-3}$. These results demonstrate that the superiority of PPLM-Contact is robust not only when using experimental structures but also when relying on predicted monomer geometries, underscoring its stability across diverse structural inputs.

Overall, the revised statistical analysis demonstrates that the performance gains of PPLM-Contact are broad, consistent, and statistically robust. Across the 18 comparisons using experimental monomer structures, 17 achieve statistical significance after FDR correction, with effect sizes ranging from medium to extremely large. When AlphaFold2-predicted structures are used, all comparisons reach significance, further reinforcing the reliability of PPLM-Contact across diverse structural conditions.

#### Text S5. Ablation analysis of PPLM-Contact

To evaluate the contribution of individual architectural modules and feature types, we trained six ablated variants of PPLM-Contact, each excluding one key component while keeping all other settings unchanged: (i) w/o cross-attn, which removes the cross-attention module from the architecture; (ii) w/o self-attn, which removes the self-attention module; (iii) w/o tri-multi, which excludes the triangle-multiplication module; (iv) w/o MSA, which excludes MSA-derived features including PSSM, DCA, and ESM-MSA-1b outputs; (v) w/o PPLM, which removes inter-protein attention features generated by PPLM; and (vi) w/o Mdist, which omits monomer distance maps extracted from either experimental or AlphaFold-predicted monomer structures.

**Figure 4D-F** illustrates the overall performance changes of these variants on the Homodimer300 and Heterodimer99 test sets. For each variant, we report the mean Top-L inter-protein contact precision and 95 % confidence intervals (CI) across three random seeds. Detailed Top-1 to Top-L results are provided in **Supplementary Table S13**. Compared with the full model (77.2 ± 0.58% for homodimers and 49.7 ± 1.02% for heterodimers), all ablations resulted in consistent and statistically meaningful declines, confirming the robustness of each component's contribution.

At the network-architecture level, removing either cross-attention or self-attention modules led to modest yet reproducible reductions—by approximately 0.8–1.0 percentage points (abbreviated as pp) for homodimers and 1.7–2.0 pp for heterodimers—indicating their supportive roles in enhancing intra- and inter-chain contextualization. In contrast, excluding the triangle-multiplication module caused a much greater decline, reducing precision by 6.7 pp and 9.4 pp on homodimers and heterodimers, respectively, highlighting its central role in capturing residue-pair geometric relationships.

At the feature level, removing the PPLM inter-protein attention features—the paired-sequence embeddings generated by PPLM—lowered precision from 77.2% to 73.3% for homodimers and from 49.7% to 45.3% for heterodimers, underscoring PPLM's key role in modeling cross-chain interactions. Removing MSA-derived features reduced precision by 7.0 pp and 9.3 pp, reflecting the contribution of co-evolutionary information, while omitting M-distance features yielded the largest degradation (23.3 pp and 16.8 pp for homodimers and heterodimers, respectively), emphasizing the indispensable role of intra-chain geometric priors. The inter-protein attention matrices contribute the smallest absolute performance gain among the three feature categories. This is expected, as we intentionally used only this single inter-protein attention feature—rather than also including sequence embeddings or intra-chain attention—to provide complementary interaction-aware information without duplicating intra-protein features that are already richly captured by structure- and MSA-based inputs.

Together, these analyses confirm that every component of PPLM-Contact contributes meaningfully to predictive accuracy, with particularly strong and statistically supported effects from the triangle-multiplication, PPLM inter-protein attention, MSA, and monomer distance modules.

#### Text S6. Analysis of the impact of MSA depth on PPLM-Contact performance

To evaluate the effect of MSA depth on model performance, we analyzed the relationship between the effective MSA depth (*Neff*) and the top-L inter-protein contact prediction precision of PPLM-Contact. Results for heterodimers (Heterodimer99 + CASP_Heterodimer20), homodimers (Homodimer300 + CASP_Homodimer43), and the combined dataset are shown in **Supplementary Figure S8**.

For heterodimers, which generally have shallower paired MSAs, precision shows a moderate positive correlation with *Neff* (Pearson's $r = 0.23$, $p = 1.1 \times 10^{-2}$). As shown in Fig. SxA–B, the average precision increases from ~30% at *Neff* < 1 to ~80% at *Neff* 30–60, indicating that PPLM-Contact can extract useful co-evolutionary information even from moderately deep alignments. Although a positive trend is observed, the explanatory power remains limited ($R^2 \approx 0.05$), suggesting that MSA depth alone does not fully account for performance variation. For homodimers, which typically have richer MSAs, no significant correlation was observed between *Neff* and precision ($r = 0.05$, $p = 0.37$; Fig. SxC–D). Precision remains consistently high across all depth bins, averaging 70–90% and reaching nearly 100% for *Neff* ⩾ 200, indicating that further MSA enrichment yields limited additional gains. When all test complexes are analyzed together (Fig. SxE–F), PPLM-Contact achieves a moderate positive correlation between Neff and precision ($r = 0.27$, $p = 2.7 \times 10^{-9}$), with precision rising from ~48% at *Neff* < 1 to ~76–91% when *Neff* > 100, albeit with large target-to-target variability ($R^2 \approx 0.07$).

The difference between heterodimers and homodimers may partly reflect how paired MSAs are constructed for these two oligomer types. For homodimers, inter-protein features are typically derived by duplicating the same single-chain MSA for both subunits, so increasing alignment depth may reinforce intra-chain rather than cross-chain coevolutionary signals, leading to performance saturation. By contrast, heterodimers rely on paired alignments of distinct proteins, where deeper MSAs can introduce additional cross-chain evolutionary constraints.

Overall, MSA depth shows a modest positive association with inter-protein contact prediction accuracy. This relationship may vary between oligomer types, with heterodimers benefiting moderately from deeper paired MSAs, whereas homodimers maintain consistently high accuracy across all depth ranges.

#### Text S7. Construction of sequence-pair datasets

We constructed a composite dataset of protein–protein interaction sequence pairs by integrating protein complex structures from the Protein Data Bank (PDB) and high-confidence interaction pairs from the STRING database. From the PDB, we extracted sequence pairs from complexes released before January 1, 2024, retaining only those in which at least one pair of residues from different chains exhibited a $C_\beta$–$C_\beta$ ($C_\alpha$ for Glycine) distance within 8Å. This yielded 273,295 heteromeric and 86,915 homomeric sequence pairs. To further expand the diversity and coverage of interaction types, we incorporated 2,995,604 high-confidence sequence pairs from STRING.

To remove redundancy at the protein-pair level, we applied a custom MMseqs2-assisted clustering procedure. For two sequence pairs $A = (A_1, A_2)$ and $B = (B_1, B_2)$, redundancy was defined only when both sequences of one pair had homologs to the two sequences of the other pair. Specifically, two pairs were considered redundant when either ($A_1$ with $B_1$ and $A_2$ with $B_2$) or ($A_1$ with $B_2$ and $A_2$ with $B_1$) satisfied at least 50% sequence identity and at least 80% alignment coverage. The identity and coverage thresholds were applied separately to each chain in the pair.

MMseqs2 was first used to identify homologs for every sequence in the dataset under the 50% identity and 80% coverage cutoffs (alignment mode 3, coverage mode 0), resulting in a homolog table for each sequence. All sequence pairs were then sorted in descending order of combined chain length. The first pair was taken as the representative of the first cluster. For each subsequent pair, we examined whether its two sequences appeared, respectively, in the homolog tables of the two sequences in any existing representative pair. If this criterion was satisfied, the pair was assigned to that cluster; otherwise, a new cluster was created with the query pair as its representative. This procedure was repeated until all pairs had been assigned.

This clustering strategy was applied separately to heteromeric PDB pairs, homomeric PDB pairs, and STRING-derived pairs, due to differences in the masking strategies used during model training. This resulted in 25,245 heteromeric clusters, 23,082 homomeric clusters, and 629,045 STRING clusters.

We then randomly selected 1,306 heteromeric, 1,194 homomeric, and 2,500 STRING-derived clusters that each contained only a single sequence pair for model validation. The remaining clusters were used for training. To ensure strict non-redundancy, we further removed all validation pairs sharing at least 50% sequence identity and at least 80% coverage with any training pair, regardless of heteromeric or homomeric status and the source of pairs. This filtering eliminated 332 pairs and yielded a final validation set of 1,204 heteromeric pairs, 1,003 homomeric pairs, and 2,471 STRING-derived pairs, for a total of 4,678 validation sequence pairs.

**Text S8. Data cleaning of PPI datasets**

The PPI datasets from D-SCRIPT include six species: *H. sapiens*, *M. musculus*, *D. melanogaster*, *C. elegans*, *S. cerevisiae*, and *E. coli*. The number of positive samples in those datasets is 421,792, 5,000, 5,000, 5,000, 5,000 and 2,000, respectively. Each dataset contains 10 times as many negative samples as positive samples, generated by randomly re-pairing proteins from different positive samples. Since a single protein can interact with diverse other proteins, the negative generation process may produce duplicate, erroneous, and invalid samples. Given proteins A and B, two samples are considered as 'duplicate' if A-B and B-A (or A-B) are generated as negative samples. A sample is defined as 'erroneous' if A-B is labeled as positive while B-A is labeled as negative. Additionally, a sample where A interacts with itself (A-A) is considered 'invalid'. **Figure S14** illustrates examples of these three cases observed in the *H. sapiens* dataset.

For duplicate samples, only one instance was retained. For erroneous data, only the positive sample was retained. All invalid samples were removed. Specifically: 362 duplicate samples, 53 erroneous samples, and 17 invalid samples was removed from *H. sapiens* dataset. 1 invalid sample was removed from *M. musculus* dataset. 2 duplicate samples and 5 invalid samples was removed from *D. melanogaster* dataset. 3 duplicate samples and 1 invalid sample was removed from *C. elegans* dataset. 28 duplicate samples, 4 erroneous samples, and 5 invalid samples was removed from *S. cerevisiae* dataset. 3765 duplicate samples and 3 invalid samples was removed from *E. coli* dataset. The final number of samples of each dataset is listed in **Table S12**.

### Supplementary Tables

**Table S1.** Performance of protein–protein interaction prediction of PPLM-PPI, TUnA, ESMDNN-PPI, D-SCRIPT, and Topsy-Turvy on the five species test sets. Bold fonts highlight the best performance in each category.

| Species | Method | Precision | Recall | Accuracy | F1-score | AUROC | AUPRC |
|---|---|---|---|---|---|---|---|
| *M. musculus* | PPLM-PPI | **0.864** | **0.843** | **0.974** | **0.853** | **0.984** | **0.920** |
| | TUnA | 0.808 | 0.819 | 0.966 | 0.814 | 0.977 | 0.885 |
| | ESMDNN-PPI | 0.781 | 0.797 | 0.961 | 0.789 | 0.975 | 0.864 |
| | D-SCRIPT | 0.817 | 0.346 | 0.934 | 0.486 | 0.833 | 0.579 |
| | Topsy-Turvy | 0.591 | 0.749 | 0.930 | 0.660 | 0.885 | 0.635 |
| *D. melanogaster* | PPLM-PPI | **0.856** | **0.818** | **0.971** | **0.837** | **0.984** | **0.906** |
| | TUnA | 0.732 | 0.813 | 0.956 | 0.770 | 0.969 | 0.853 |
| | ESMDNN-PPI | 0.737 | 0.794 | 0.956 | 0.765 | 0.969 | 0.843 |
| | D-SCRIPT | 0.799 | 0.359 | 0.934 | 0.496 | 0.824 | 0.553 |
| | Topsy-Turvy | 0.621 | 0.741 | 0.935 | 0.676 | 0.896 | 0.649 |
| *C. elegans* | PPLM-PPI | **0.914** | 0.696 | **0.966** | **0.790** | **0.976** | **0.883** |
| | TUnA | 0.786 | **0.724** | 0.957 | 0.754 | 0.957 | 0.824 |
| | ESMDNN-PPI | 0.788 | 0.681 | 0.954 | 0.731 | 0.955 | 0.799 |
| | D-SCRIPT | 0.840 | 0.305 | 0.932 | 0.448 | 0.813 | 0.548 |
| | Topsy-Turvy | 0.682 | 0.645 | 0.940 | 0.663 | 0.851 | 0.616 |
| *S. cerevisiae* | PPLM-PPI | **0.722** | **0.652** | **0.945** | **0.685** | **0.935** | **0.745** |
| | TUnA | 0.536 | 0.647 | 0.917 | 0.586 | 0.896 | 0.633 |
| | ESMDNN-PPI | 0.565 | 0.637 | 0.922 | 0.599 | 0.919 | 0.619 |
| | D-SCRIPT | 0.708 | 0.223 | 0.921 | 0.339 | 0.789 | 0.405 |
| | Topsy-Turvy | 0.484 | 0.544 | 0.906 | 0.512 | 0.762 | 0.432 |
| *E. coli* | PPLM-PPI | 0.725 | **0.709** | **0.939** | **0.717** | **0.910** | **0.784** |
| | TUnA | 0.581 | 0.666 | 0.911 | 0.621 | 0.887 | 0.694 |
| | ESMDNN-PPI | 0.680 | 0.601 | 0.925 | 0.638 | 0.899 | 0.685 |
| | D-SCRIPT | **0.790** | 0.383 | 0.921 | 0.516 | 0.860 | 0.574 |
| | Topsy-Turvy | 0.451 | 0.464 | 0.879 | 0.458 | 0.643 | 0.406 |

**Table S2.** Mean and standard deviation of performance metrics for PPLM-PPI and comparison methods across the five species test sets.

| Method | Precision | Recall | Accuracy | F1-score | AUROC | AUPRC |
|---|---|---|---|---|---|---|
| PPLM-PPI | 0.816±0.087 | 0.744±0.083 | 0.959±0.016 | 0.776±0.073 | 0.958±0.034 | 0.848±0.078 |
| TUnA | 0.689±0.123 | 0.734±0.080 | 0.941±0.025 | 0.709±0.100 | 0.937±0.043 | 0.778±0.109 |
| ESMDNN-PPI | 0.710±0.092 | 0.702±0.090 | 0.944±0.019 | 0.704±0.082 | 0.943±0.033 | 0.762±0.106 |
| D-SCRIPT | 0.791±0.050 | 0.323±0.063 | 0.928±0.007 | 0.457±0.070 | 0.824±0.026 | 0.532±0.072 |
| Topsy-Turvy | 0.566±0.096 | 0.629±0.124 | 0.918±0.025 | 0.594±0.101 | 0.807±0.106 | 0.548±0.118 |

**Table S3.** Statistical comparison between PPLM-PPI and baseline models (TUnA, ESMDNN-PPI, D-SCRIPT, and Topsy-Turvy) across the five species test sets. For each evaluation metric (F1-score, AUROC, and AUPRC), the table reports the fold-wise mean difference (Δ), relative percentage improvement, and paired effect size (Cohen's d).

| | Metric | Mean Δ | Percent gain (%) | Cohen's d |
|---|---|---|---|---|
| ***vs TUnA*** | F1-score | 0.068 | 10.1 | 2.252 |
| | AUROC | 0.021 | 2.3 | 1.742 |
| | AUPRC | 0.070 | 9.6 | 2.268 |
| ***vs ESMDNN-PPI*** | F1-score | 0.072 | 10.5 | 6.586 |
| | AUROC | 0.014 | 1.5 | 3.084 |
| | AUPRC | 0.086 | 11.9 | 3.026 |
| ***vs D-SCRIPT*** | F1-score | 0.319 | 72.3 | 4.765 |
| | AUROC | 0.134 | 16.4 | 2.824 |
| | AUPRC | 0.316 | 60.9 | 5.306 |
| ***vs Topsy-Turvy*** | F1-score | 0.183 | 32.5 | 3.728 |
| | AUROC | 0.15 | 20.0 | 2.062 |
| | AUPRC | 0.3 | 58.7 | 6.183 |

**Table S4.** Performance of PPLM-PPI ablation variants using different pooling strategies across five species test sets. Values represent the mean accuracy ± 95 % confidence intervals (CI) averaged over three random seeds.

| Method | Precision | Recall | Accuracy | F1-score | AUROC | AUPRC |
|---|---|---|---|---|---|---|
| ***M. musculus*** | | | | | | |
| mean_max | **0.875±0.011** | **0.840±0.005** | **0.974±0.001** | **0.857±0.004** | **0.984±0**.00 | **0.921±0.002** |
| mean_max_min | 0.881±0.008 | 0.817±0.003 | 0.973±0.001 | 0.848±0.002 | 0.983±0.00 | 0.916±0.001 |
| mean_min | 0.876±0.011 | 0.808±0.008 | 0.972±0.00 | 0.841±0.001 | 0.982±0.00 | 0.908±0.001 |
| max_min | 0.867±0.002 | 0.792±0.005 | 0.970±0.00 | 0.828±0.002 | 0.980±0.00 | 0.901±0.001 |
| mean | 0.854±0.025 | 0.838±0.019 | 0.972±0.001 | 0.845±0.003 | 0.983±0.00 | 0.914±0.002 |
| max | 0.851±0.003 | 0.824±0.005 | 0.971±0.001 | 0.837±0.003 | 0.980±0.00 | 0.903±0.002 |
| ***D. melanogaster*** | | | | | | |
| mean_max | **0.864±0.008** | **0.807±0.012** | **0.971±0.00** | **0.834±0.003** | **0.984±0.00** | **0.906±0.002** |
| mean_max_min | 0.877±0.002 | 0.781±0.002 | 0.970±0.00 | 0.827±0.002 | 0.983±0.00 | 0.900±0.001 |
| mean_min | 0.865±0.005 | 0.768±0.009 | 0.968±0.001 | 0.813±0.004 | 0.979±0.001 | 0.885±0.001 |
| max_min | 0.865±0.005 | 0.745±0.013 | 0.966±0.001 | 0.801±0.007 | 0.979±0.001 | 0.878±0.004 |
| mean | 0.816±0.029 | 0.808±0.022 | 0.966±0.002 | 0.811±0.005 | 0.978±0.001 | 0.886±0.005 |
| max | 0.842±0.004 | 0.779±0.009 | 0.967±0.001 | 0.809±0.006 | 0.979±0.001 | 0.880±0.007 |
| ***C. elegans*** | | | | | | |
| mean_max | **0.918±0.004** | **0.684±0.012** | **0.966±0.001** | **0.784±0.007** | **0.976±0.001** | **0.884±0.002** |
| mean_max_min | 0.929±0.004 | 0.651±0.004 | 0.964±0.00 | 0.766±0.002 | 0.976±0.001 | 0.881±0.003 |
| mean_min | 0.907±0.008 | 0.643±0.012 | 0.962±0.001 | 0.752±0.007 | 0.971±0.001 | 0.860±0.002 |
| max_min | 0.921±0.010 | 0.607±0.015 | 0.959±0.001 | 0.731±0.008 | 0.973±0.002 | 0.862±0.005 |
| mean | 0.859±0.022 | 0.703±0.031 | 0.962±0.000 | 0.773±0.010 | 0.967±0.001 | 0.853±0.003 |
| max | 0.903±0.005 | 0.654±0.006 | 0.962±0.001 | 0.758±0.004 | 0.973±0.002 | 0.863±0.006 |
| ***S. cerevisiae*** | | | | | | |
| **mean_max** | **0.741±0.019** | **0.638±0.014** | **0.947±0.001** | **0.685±0.001** | **0.935±0.001** | **0.745±0.00** |
| mean_max_min | 0.756±0.009 | 0.618±0.008 | 0.947±0.000 | 0.680±0.001 | 0.934±0.001 | 0.738±0.003 |
| mean_min | 0.722±0.022 | 0.625±0.014 | 0.944±0.002 | 0.669±0.002 | 0.929±0.001 | 0.721±0.005 |
| max_min | 0.736±0.007 | 0.582±0.010 | 0.943±0.00 | 0.650±0.004 | 0.925±0.002 | 0.707±0.005 |
| mean | 0.660±0.036 | 0.669±0.032 | 0.938±0.004 | 0.664±0.005 | 0.929±0.001 | 0.723±0.003 |
| max | 0.707±0.012 | 0.601±0.006 | 0.941±0.001 | 0.650±0.003 | 0.926±0.00 | 0.707±0.002 |
| ***E. coli*** | | | | | | |
| mean_max | **0.726±0.021** | **0.701±0.015** | **0.938±0.002** | **0.713±0.005** | **0.909±0.001** | **0.782±0.002** |
| mean_max_min | 0.741±0.010 | 0.689±0.004 | 0.940±0.001 | 0.714±0.005 | 0.912±0.003 | 0.784±0.002 |
| mean_min | 0.661±0.011 | 0.697±0.003 | 0.928±0.002 | 0.679±0.006 | 0.906±0.004 | 0.769±0.004 |
| max_min | 0.708±0.007 | 0.670±0.002 | 0.933±0.001 | 0.688±0.004 | 0.906±0.003 | 0.767±0.003 |
| mean | 0.591±0.036 | 0.707±0.014 | 0.914±0.008 | 0.643±0.017 | 0.902±0.002 | 0.754±0.007 |
| max | 0.686±0.057 | 0.673±0.023 | 0.930±0.007 | 0.678±0.015 | 0.901±0.003 | 0.757±0.004 |

**Table S5.** Performance of PPLM-PPI ablation variants using different features across five species test sets. Values represent the mean accuracy ± 95 % confidence intervals (CI) averaged over three random seeds.

| Method | Precision | Recall | Accuracy | F1-score | AUROC | AUPRC |
|---|---|---|---|---|---|---|
| ***M. musculus*** | | | | | | |
| inter-attn + intra-attn + embed | **0.854±0.025** | **0.838±0.019** | **0.972±0.001** | **0.845±0.003** | **0.983±0.000** | **0.914±0.002** |
| inter_attn + intra_attn | 0.742±0.044 | 0.315±0.034 | 0.928±0.001 | 0.441±0.024 | 0.866±0.000 | 0.560±0.001 |
| intra_attn + embed | 0.847±0.014 | 0.795±0.002 | 0.968±0.000 | 0.820±0.005 | 0.979±0.000 | 0.890±0.003 |
| embed | 0.837±0.011 | 0.796±0.014 | 0.967±0.000 | 0.816±0.003 | 0.978±0.001 | 0.887±0.004 |
| intra_attn | 0.415±0.063 | 0.591±0.061 | 0.885±0.021 | 0.484±0.021 | 0.853±0.001 | 0.525±0.003 |
| ***D. melanogaster*** | | | | | | |
| inter-attn + intra-attn + embed | **0.816±0.029** | **0.808±0.022** | **0.966±0.002** | **0.811±0.005** | **0.978±0.001** | **0.886±0.005** |
| inter_attn + intra_attn | 0.638±0.040 | 0.441±0.045 | 0.926±0.003 | 0.520±0.016 | 0.903±0.002 | 0.586±0.007 |
| intra_attn + embed | 0.798±0.032 | 0.735±0.024 | 0.959±0.002 | 0.765±0.005 | 0.969±0.002 | 0.843±0.007 |
| embed | 0.786±0.020 | 0.733±0.012 | 0.958±0.002 | 0.758±0.005 | 0.968±0.001 | 0.837±0.001 |
| intra_attn | 0.470±0.070 | 0.740±0.056 | 0.898±0.021 | 0.571±0.034 | 0.915±0.001 | 0.659±0.005 |
| ***C. elegans*** | | | | | | |
| inter-attn + intra-attn + embed | **0.859±0.022** | **0.703±0.031** | **0.962±0.000** | **0.773±0.010** | **0.967±0.001** | **0.853±0.003** |
| inter_attn + intra_attn | 0.814±0.036 | 0.331±0.039 | 0.932±0.001 | 0.470±0.033 | 0.923±0.000 | 0.657±0.004 |
| intra_attn + embed | 0.848±0.033 | 0.643±0.016 | 0.957±0.002 | 0.731±0.003 | 0.959±0.005 | 0.819±0.014 |
| embed | 0.803±0.016 | 0.644±0.008 | 0.956±0.001 | 0.725±0.002 | 0.959±0.003 | 0.812±0.007 |
| intra_attn | 0.615±0.075 | 0.648±0.075 | 0.930±0.009 | 0.626±0.004 | 0.930±0.001 | 0.692±0.004 |
| ***S. cerevisiae*** | | | | | | |
| inter-attn + intra-attn + embed | 0.66±0.036 | **0.669±0.032** | 0.938±0.004 | **0.664±0.005** | **0.929±0.001** | **0.723±0.003** |
| inter_attn + intra_attn | 0.601±0.039 | 0.372±0.043 | 0.920±0.003 | 0.457±0.020 | 0.879±0.000 | 0.522±0.000 |
| intra_attn + embed | **0.689±0.054** | 0.609±0.040 | **0.939±0.005** | 0.645±0.007 | 0.924±0.003 | 0.701±0.009 |
| embed | 0.619±0.036 | 0.641±0.005 | 0.931±0.005 | 0.630±0.016 | 0.921±0.004 | 0.687±0.012 |
| intra_attn | 0.336±0.047 | 0.668±0.075 | 0.847±0.031 | 0.444±0.024 | 0.871±0.002 | 0.502±0.008 |
| ***E. coli*** | | | | | | |
| inter-attn + intra-attn + embed | 0.591±0.036 | **0.707±0.014** | 0.914±0.008 | **0.643±0.017** | **0.902±0.002** | **0.754±0.007** |
| inter_attn + intra_attn | **0.883±0.020** | 0.492±0.008 | **0.937±0.001** | 0.631±0.004 | 0.801±0.005 | 0.623±0.005 |
| intra_attn + embed | 0.601±0.065 | 0.679±0.033 | 0.914±0.013 | 0.636±0.024 | 0.893±0.003 | 0.736±0.001 |
| embed | 0.603±0.067 | 0.687±0.026 | 0.915±0.012 | 0.641±0.028 | 0.888±0.007 | 0.731±0.011 |
| intra_attn | 0.329±0.061 | 0.644±0.047 | 0.812±0.045 | 0.432±0.042 | 0.807±0.002 | 0.613±0.003 |

**Table S6.** AUPRC of PPLM-PPI and baseline methods across sequence-identity bins, defined by the maximum single-sequence identity of either protein to any human protein in the training set, for all five species.

| Method | (0, 0.5] | (0.5, 0.6] | (0.6, 0.7] | (0.7, 0.8] | (0.8, 0.9] | (0.9, 1.0) | [1.0] |
|---|---|---|---|---|---|---|---|
| ***M. musculus*** | | | | | | | |
| Samples (n) | 1398 | 876 | 1654 | 4778 | 13246 | 30191 | 2856 |
| Neg : Pos ratio | 86.4 | 61.6 | 30.2 | 19.8 | 14.3 | 9.8 | 1.8 |
| PPLM-PPI | 0.605 | 0.842 | 0.885 | 0.929 | 0.930 | 0.895 | 0.974 |
| TUnA | 0.480 | 0.866 | 0.777 | 0.880 | 0.883 | 0.854 | 0.964 |
| ESMDNN-PPI | 0.464 | 0.866 | 0.781 | 0.864 | 0.867 | 0.819 | 0.962 |
| D-SCRIPT | 0.142 | 0.370 | 0.390 | 0.502 | 0.623 | 0.479 | 0.856 |
| Topsy-Turvy | 0.124 | 0.233 | 0.337 | 0.538 | 0.639 | 0.572 | 0.867 |
| ***D. melanogaster*** | | | | | | | |
| Samples (n) | 38443 | 5464 | 4470 | 3220 | 2201 | 1053 | 142 |
| Neg : Pos ratio | 22.2 | 8.0 | 4.5 | 2.7 | 1.9 | 2.8 | 46.3 |
| PPLM-PPI | 0.859 | 0.877 | 0.930 | 0.950 | 0.949 | 0.896 | 0.817 |
| TUnA | 0.762 | 0.822 | 0.891 | 0.924 | 0.938 | 0.885 | 0.333 |
| ESMDNN-PPI | 0.763 | 0.823 | 0.866 | 0.907 | 0.917 | 0.879 | 0.903 |
| D-SCRIPT | 0.401 | 0.448 | 0.657 | 0.755 | 0.811 | 0.646 | 0.014 |
| Topsy-Turvy | 0.480 | 0.622 | 0.741 | 0.782 | 0.846 | 0.695 | 0.372 |
| ***C. elegans*** | | | | | | | |
| Samples (n) | 44951 | 3728 | 2736 | 1865 | 1275 | 441 | 0 |
| Neg : Pos ratio | 21.4 | 4.1 | 2.4 | 1.6 | 1.1 | 2.0 | NA |
| PPLM-PPI | 0.748 | 0.910 | 0.950 | 0.972 | 0.978 | 0.882 | NA |
| TUnA | 0.657 | 0.873 | 0.911 | 0.958 | 0.953 | 0.844 | NA |
| ESMDNN-PPI | 0.629 | 0.835 | 0.911 | 0.913 | 0.961 | 0.858 | NA |
| D-SCRIPT | 0.308 | 0.521 | 0.789 | 0.825 | 0.845 | 0.690 | NA |
| Topsy-Turvy | 0.372 | 0.698 | 0.812 | 0.882 | 0.893 | 0.734 | NA |
| ***S. cerevisiae*** | | | | | | | |
| Samples (n) | 45865 | 4023 | 2915 | 1552 | 478 | 130 | 0 |
| Neg : Pos ratio | 15.6 | 4.7 | 2.8 | 1.8 | 1.8 | 1.7 | NA |
| PPLM-PPI | 0.659 | 0.769 | 0.850 | 0.907 | 0.874 | 0.888 | NA |
| TUnA | 0.494 | 0.709 | 0.802 | 0.866 | 0.812 | 0.805 | NA |
| ESMDNN-PPI | 0.486 | 0.669 | 0.802 | 0.843 | 0.829 | 0.848 | NA |
| D-SCRIPT | 0.240 | 0.488 | 0.665 | 0.773 | 0.747 | 0.684 | NA |
| Topsy-Turvy | 0.294 | 0.553 | 0.642 | 0.732 | 0.710 | 0.694 | NA |
| ***E. coli*** | | | | | | | |
| Samples (n) | 17857 | 257 | 100 | 18 | 0 | 0 | 0 |
| Neg : Pos ratio | 8.5 | 2.3 | 2.3 | 2.0 | NA | NA | NA |
| PPLM-PPI | 0.777 | 0.890 | 0.862 | 0.974 | NA | NA | NA |
| TUnA | 0.686 | 0.815 | 0.709 | 0.944 | NA | NA | NA |
| ESMDNN-PPI | 0.678 | 0.773 | 0.783 | 0.910 | NA | NA | NA |
| D-SCRIPT | 0.565 | 0.790 | 0.701 | 0.386 | NA | NA | NA |
| Topsy-Turvy | 0.383 | 0.757 | 0.720 | 0.784 | NA | NA | NA |

**Table S7.** Fold-wise performance of PPLM-Affinity, ESM2-Affinity, and PPB-Affinity across the five cross-validation splits. Performance metrics (PCC, SRCC, and RMSD) are reported separately for each fold.

| #CV | Method | Entire dataset | | | Antibody–antigen subgroup | | | TCR–pMHC subgroup | | |
|---|---|---|---|---|---|---|---|---|---|---|
| | | PCC | SRCC | RMSD | PCC | SRCC | RMSD | PCC | SRCC | RMSD |
| CV 0 | PPLM-Affinity | 0.676 | 0.743 | 2.716 | 0.548 | 0.526 | 1.802 | 0.131 | 0.225 | 1.233 |
| | ESM2-Affinity | 0.504 | 0.526 | 3.130 | 0.178 | 0.140 | 2.038 | 0.196 | 0.152 | 1.734 |
| | PPB-Affinity | 0.492 | 0.549 | 3.144 | 0.487 | 0.486 | 1.761 | 0.150 | 0.140 | 1.846 |
| CV 1 | PPLM-Affinity | 0.710 | 0.690 | 2.062 | 0.384 | 0.412 | 1.794 | 0.327 | 0.327 | 1.875 |
| | ESM2-Affinity | 0.558 | 0.570 | 2.388 | 0.231 | 0.298 | 1.755 | 0.233 | 0.167 | 1.684 |
| | PPB-Affinity | 0.668 | 0.677 | 2.119 | 0.238 | 0.279 | 1.913 | 0.286 | 0.335 | 1.693 |
| CV 2 | PPLM-Affinity | 0.624 | 0.610 | 2.191 | 0.437 | 0.454 | 1.642 | 0.530 | 0.249 | 1.852 |
| | ESM2-Affinity | 0.509 | 0.497 | 2.342 | 0.165 | 0.141 | 1.781 | 0.007 | -0.139 | 2.755 |
| | PPB-Affinity | 0.509 | 0.491 | 2.382 | 0.114 | 0.088 | 1.938 | 0.531 | 0.412 | 1.684 |
| CV 3 | PPLM-Affinity | 0.556 | 0.532 | 2.532 | 0.351 | 0.386 | 2.088 | 0.405 | 0.481 | 1.688 |
| | ESM2-Affinity | 0.518 | 0.512 | 2.357 | 0.116 | 0.230 | 2.401 | 0.392 | 0.349 | 2.714 |
| | PPB-Affinity | 0.552 | 0.541 | 2.320 | 0.171 | 0.258 | 2.120 | -0.158 | -0.173 | 2.319 |
| CV4 | PPLM-Affinity | 0.647 | 0.607 | 2.058 | 0.178 | 0.243 | 2.206 | 0.435 | 0.277 | 1.519 |
| | ESM2-Affinity | 0.649 | 0.629 | 2.161 | 0.184 | 0.148 | 2.243 | 0.472 | 0.155 | 1.603 |
| | PPB-Affinity | 0.506 | 0.442 | 2.352 | 0.175 | 0.176 | 2.252 | 0.506 | 0.426 | 1.669 |

**Table S8.** Mean performance of PPLM-Affinity, ESM2-Affinity, and PPB-Affinity on the full affinity dataset, the antibody–antigen subset, and the TCR–pMHC subset across five-fold cross-validation. Values represent mean ± standard deviation of PCC, SRCC, and RMSE computed over the five folds.

| Dataset | Method | PCC (mean ± s.d.) | SRCC (mean ± s.d.) | RMSE (mean ± s.d.) |
|---|---|---|---|---|
| Entire dataset | PPLM-Affinity | 0.643±0.058 | 0.636±0.082 | 2.312±0.297 |
| | ESM2-Affinity | 0.548±0.061 | 0.547±0.053 | 2.476±0.376 |
| | PPB-Affinity | 0.545±0.072 | 0.540±0.088 | 2.463±0.394 |
| Antibody–antigen subgroup | PPLM-Affinity | 0.380±0.135 | 0.404±0.105 | 1.906±0.232 |
| | ESM2-Affinity | 0.175±0.041 | 0.191±0.071 | 2.044±0.283 |
| | PPB-Affinity | 0.237±0.146 | 0.257±0.148 | 1.997±0.191 |
| TCR–pMHC subgroup | PPLM-Affinity | 0.366±0.150 | 0.312±0.102 | 1.633±0.266 |
| | ESM2-Affinity | 0.260±0.181 | 0.137±0.175 | 2.098±0.583 |
| | PPB-Affinity | 0.263±0.283 | 0.228±0.252 | 1.842±0.276 |

**Table S9.** Statistical comparison between PPLM-Affinity and baseline models (ESM2-Affinity and PPB-Affinity) across five-fold cross-validation. For each dataset and metric, the table reports the fold-wise mean difference (Δ), relative percentage improvement, and paired effect size (Cohen's d).

| **Dataset** | **PCC** | | | **SRCC** | | | **RMSE** | | |
|---|---|---|---|---|---|---|---|---|---|
| | **Mean Δ** | **Percent gain (%)** | **Cohen's d** | **Mean Δ** | **Percent gain (%)** | **Cohen's d** | **Mean Δ** | **Percent gain (%)** | **Cohen's d** |
| ***vs ESM2-Affinity*** | | | | | | | | | |
| Entire dataset | 0.095 | 17.3 | 1.274 | 0.09 | 16.4 | 0.958 | 0.164 | 6.6 | 0.719 |
| Antibody–antigen subgroup | 0.205 | 117.1 | 1.45 | 0.213 | 111.5 | 1.645 | 0.137 | 6.7 | 0.96 |
| TCR–pMHC subgroup | 0.216 | 144.0 | 0.706 | 0.175 | 127.7 | 1.421 | 0.465 | 22.2 | 0.893 |
| ***vs PPB-Affinity*** | | | | | | | | | |
| Entire dataset | 0.097 | 18.0 | 1.326 | 0.096 | 17.8 | 1.064 | 0.152 | 6.2 | 0.62 |
| Antibody–antigen subgroup | 0.143 | 60.3 | 1.163 | 0.147 | 57.2 | 1.14 | 0.09 | 4.5 | 0.705 |
| TCR–pMHC subgroup | 0.103 | 39.2 | 0.394 | 0.084 | 36.8 | 0.25 | 0.209 | 11.4 | 0.522 |

**Table S10.** Precision (%) of inter-protein contact prediction for PPLM-Contact, PLMGraph-Inter, DeepInter, CDPred, DeepHomo2.0, and GLINTER on the Homodimer300, Heterodimer99, CASP_Homo43, and CASP_Hetero20 test sets, using both experimental and AlphaFold2-predicted (values in parentheses) monomer structures. $L$ denotes the length of the shorter protein in the dimer.

| **Method** | **Top 1** | **Top 10** | **Top 50** | **Top *L*/10** | **Top *L*/5** | **Top *L*** |
|---|---|---|---|---|---|---|
| ***Homodimer300 test set*** | | | | | | |
| PPLM-Contact | 86.0 (75.7) | 84.5 (73.7) | 82.4 (72.1) | 83.6 (72.8) | 82.5 (72.4) | 77.8 (66.6) |
| PLMGraph-Inter | 71.7 (66.0) | 69.4 (61.5) | 62.6 (55.0) | 67.2 (58.9) | 63.7 (56.0) | 50.6 (44.1) |
| DeepInter | 76.3 (70.3) | 75.2 (68.5) | 73.7 (66.1) | 74.8 (67.6) | 74.3 (66.7) | 68.9 (60.3) |
| CDPred | 77.0 (70.0) | 76.0 (68.3) | 72.3 (65.0) | 74.4 (67.0) | 72.4 (65.3) | 63.0 (56.0) |
| DeepHomo2.0 | 68.7 (59.7) | 67.1 (57.6) | 63.2 (54.4) | 66.0 (56.8) | 63.5 (54.6) | 51.4 (44.2) |
| GLINTER | 53.7 (47.0) | 51.1 (46.7) | 44.5 (39.9) | 48.9 (44.9) | 46.3 (41.3) | 34.5 (30.7) |
| ***Heterodimer99 test set*** | | | | | | |
| PPLM-Contact | 62.6 (56.6) | 58.1 (54.0) | 54.9 (50.1) | 57.0 (53.6) | 56.5 (52.4) | 48.9 (45.1) |
| PLMGraph-Inter | 49.5 (38.4) | 46.2 (36.3) | 37.3 (29.3) | 43.3 (35.2) | 41.1 (31.9) | 31.0 (25.0) |
| DeepInter | 40.4 (42.4) | 41.4 (38.2) | 38.6 (36.5) | 40.7 (38.6) | 39.7 (37.0) | 35.0 (32.9) |
| CDPred | 34.3 (37.4) | 30.9 (37.5) | 28.5 (31.5) | 31.4 (37.9) | 30.0 (35.5) | 25.1 (27.5) |
| GLINTER | 27.3 (26.3) | 25.9 (24.6) | 21.2 (20.5) | 26.4 (24.5) | 23.4 (22.4) | 17.3 (16.8) |
| ***CASP_ Homodimer43 test set*** | | | | | | |
| PPLM-Contact | 69.8 (67.4) | 70.9 (63.7) | 69.2 (59.7) | 71.5 (64.3) | 70.7 (61.0) | 65.2 (54.0) |
| PLMGraph-Inter | 66.7 (57.1) | 65.5 (54.8) | 57.3 (48.3) | 63.3 (53.3) | 60.3 (50.8) | 46.1 (39.6) |
| DeepInter | 69.8 (59.3) | 69.1 (54.8) | 67.3 (51.9) | 69.2 (54.1) | 68.1 (52.6) | 61.5 (47.8) |
| CDPred | 55.8 (48.8) | 57.2 (46.3) | 50.9 (41.7) | 54.1 (45.6) | 52.5 (42.5) | 44.4 (36.9) |
| DeepHomo2.0 | 50.0 (47.6) | 46.4 (43.1) | 42.1 (38.9) | 44.4 (44.0) | 43.5 (40.0) | 33.4 (31.3) |
| GLINTER | 44.2 (39.5) | 42.3 (34.4) | 36.3 (30.1) | 40.7 (31.8) | 36.2 (30.8) | 25.3 (22.5) |
| ***CASP_ Heterodimer20 test set*** | | | | | | |
| PPLM-Contact | 60.0 (50.0) | 54.0 (51.0) | 49.4 (45.1) | 53.0 (49.0) | 50.8 (45.8) | 45.6 (40.5) |
| PLMGraph-Inter | 40.0 (35.0) | 34.0 (33.5) | 27.2 (26.4) | 32.5 (33.1) | 30.9 (29.1) | 23.0 (23.3) |
| DeepInter | 25.0 (40.0) | 32.5 (37.5) | 29.7 (33.0) | 33.0 (36.4) | 30.9 (35.5) | 26.7 (29.7) |
| CDPred | 30.0 (25.0) | 31.0 (27.0) | 24.5 (21.8) | 33.0 (27.4) | 30.4 (25.9) | 20.4 (19.7) |
| GLINTER | 20.0 (20.0) | 21.0 (12.5) | 12.5 (7.3) | 20.0 (11.6) | 16.7 (10.3) | 10.3 (6.9) |

**Table S11.** Comprehensive statistical comparison between PPLM-Contact and baseline models (PLMGraph-Inter, DeepInter, CDPred, DeepHomo2.0, and GLINTER) across four test datasets (Homodimer300, Heterodimer99, CASP_Homo43, and CASP_Hetero20) using distance maps derived from experimental monomer structures. For each dataset and baseline, improvements in per-target top *L* contact precision were evaluated using mean difference (Δ), relative percent gain, paired effect size (Cohen's d), Wilcoxon signed-rank *p*-values, and Benjamini–Hochberg FDR-adjusted *q*-values. Statistically significant results ($q < 0.05$) are indicated.

| Dataset | Mean Δ | Percent gain (%) | Effect size (Cohen's d) | Wilcoxon p-value | FDR q-value | Significant (q < 0.05) |
|---|---|---|---|---|---|---|
| ***vs PLMGraph-Inter*** | | | | | | |
| Homodimer300 | 27.19 | 53.7 | 1.04 | 1.06E-38 | 4.23E-38 | ✓ |
| Heterodimer99 | 17.94 | 57.9 | 0.57 | 1.14E-06 | 2.28E-06 | ✓ |
| CASP_Homodimer43 | 20.20 | 44.8 | 0.59 | 3.75E-04 | 5.00E-04 | ✓ |
| CASP_Heterodimer20 | 22.57 | 98.1 | 0.68 | 9.56E-03 | 9.56E-03 | ✓ |
| ***vs DeepInter*** | | | | | | |
| Homodimer300 | 8.85 | 12.8 | 0.44 | 3.63E-21 | 1.45E-20 | ✓ |
| Heterodimer99 | 13.89 | 39.7 | 0.43 | 3.05E-05 | 6.10E-05 | ✓ |
| CASP_Homodimer43 | 3.71 | 6.0 | 0.15 | 7.42E-02 | 7.42E-02 | × |
| CASP_Heterodimer20 | 18.86 | 70.6 | 0.95 | 2.93E-04 | 3.91E-04 | ✓ |
| ***vs CDPred*** | | | | | | |
| Homodimer300 | 14.82 | 23.5 | 0.64 | 5.16E-31 | 2.06E-30 | ✓ |
| Heterodimer99 | 23.83 | 94.9 | 0.69 | 1.03E-08 | 2.05E-08 | ✓ |
| CASP_Homodimer43 | 20.80 | 46.8 | 0.57 | 4.97E-04 | 6.63E-04 | ✓ |
| CASP_Heterodimer20 | 25.13 | 123.1 | 0.87 | 3.28E-03 | 3.28E-03 | ✓ |
| ***vs DeepHomo2.0*** | | | | | | |
| Homodimer300 | 26.38 | 51.3 | 1.08 | 3.11E-43 | 6.22E-43 | ✓ |
| CASP_Homodimer43 | 31.80 | 95.1 | 1.06 | 1.96E-07 | 1.96E-07 | ✓ |
| ***vs GLINTER*** | | | | | | |
| Homodimer300 | 43.26 | 125.3 | 1.59 | 1.29E-47 | 5.18E-47 | ✓ |
| Heterodimer99 | 31.63 | 182.7 | 0.96 | 1.15E-12 | 2.29E-12 | ✓ |
| CASP_Homodimer43 | 39.90 | 157.7 | 1.21 | 2.42E-07 | 3.22E-07 | ✓ |
| CASP_Heterodimer20 | 35.23 | 341.4 | 1.11 | 3.42E-04 | 3.42E-04 | ✓ |

**Table S12.** Comprehensive statistical comparison between PPLM-Contact and baseline models (PLMGraph-Inter, DeepInter, CDPred, DeepHomo2.0, and GLINTER) across four test datasets (Homodimer300, Heterodimer99, CASP_Homo43, and CASP_Hetero20) using distance maps derived from AlphaFold2-predicted monomer structures. For each dataset and baseline, improvements in per-target top *L* contact precision were evaluated using mean difference (Δ), relative percent gain, paired effect size (Cohen's d), Wilcoxon signed-rank *p*-values, and Benjamini–Hochberg FDR-adjusted *q*-values. Statistically significant results ($q < 0.05$) are indicated.

| Dataset | Mean Δ | Percent gain (%) | Effect size (Cohen's d) | Wilcoxon p-value | FDR q-value | Significant (q < 0.05) |
|---|---|---|---|---|---|---|
| ***vs PLMGraph-Inter*** | | | | | | |
| Homodimer300 | 22.52 | 51.1 | 0.88 | 3.92E-33 | 1.57E-32 | ✓ |
| Heterodimer99 | 20.07 | 80.2 | 0.67 | 2.39E-08 | 4.77E-08 | ✓ |
| CASP_Homodimer43 | 15.29 | 39.5 | 0.51 | 1.53E-03 | 2.04E-03 | ✓ |
| CASP_Heterodimer20 | 17.20 | 73.9 | 0.53 | 4.95E-02 | 4.95E-02 | ✓ |
| ***vs DeepInter*** | | | | | | |
| Homodimer300 | 6.27 | 10.4 | 0.35 | 3.96E-13 | 1.59E-12 | ✓ |
| Heterodimer99 | 12.18 | 37.0 | 0.43 | 6.10E-05 | 1.22E-04 | ✓ |
| CASP_Homodimer43 | 6.17 | 12.9 | 0.27 | 1.30E-02 | 1.30E-02 | ✓ |
| CASP_Heterodimer20 | 10.78 | 36.3 | 0.68 | 4.29E-03 | 5.71E-03 | ✓ |
| ***vs CDPred*** | | | | | | |
| Homodimer300 | 10.54 | 18.8 | 0.48 | 3.43E-21 | 1.37E-20 | ✓ |
| Heterodimer99 | 17.64 | 64.2 | 0.55 | 1.69E-06 | 3.38E-06 | ✓ |
| CASP_Homodimer43 | 17.09 | 46.4 | 0.66 | 1.51E-04 | 2.02E-04 | ✓ |
| CASP_Heterodimer20 | 20.81 | 105.9 | 0.65 | 2.00E-02 | 2.00E-02 | ✓ |
| ***vs DeepHomo2.0*** | | | | | | |
| Homodimer300 | 22.43 | 50.8 | 0.89 | 1.68E-37 | 3.37E-37 | ✓ |
| CASP_Homodimer43 | 22.63 | 72.3 | 0.84 | 1.02E-05 | 1.02E-05 | ✓ |
| ***vs GLINTER*** | | | | | | |
| Homodimer300 | 35.85 | 116.7 | 1.25 | 1.30E-42 | 5.19E-42 | ✓ |
| Heterodimer99 | 28.28 | 167.9 | 0.88 | 1.29E-11 | 2.58E-11 | ✓ |
| CASP_Homodimer43 | 31.48 | 140.0 | 1.09 | 5.33E-07 | 7.11E-07 | ✓ |
| CASP_Heterodimer20 | 33.52 | 483.3 | 0.94 | 2.26E-03 | 2.26E-03 | ✓ |

**Table S13.** Performance of PPLM-Contact and its ablation variants on the Homodimer300 and Heterodimer99 test sets. Values indicate the mean Top-L inter-protein contact precision (%) ± 95 % confidence intervals (CI) computed across three random seeds.

| Method | Top 1 | Top 10 | Top 50 | Top *L*/10 | Top *L*/5 | Top *L* |
|---|---|---|---|---|---|---|
| ***Homodimer300 test set*** | | | | | | |
| PPLM-Contact | 85.3±0.75 | 84.1±0.46 | 82.0±0.50 | 83.2±0.48 | 82.2±0.38 | 77.2±0.58 |
| w/o cross-attn | 85.0±0.75 | 83.3±0.38 | 81.2±0.38 | 82.4±0.10 | 81.5±0.18 | 76.2±0.29 |
| w/o self-attn | 84.7±0.75 | 83.8±0.19 | 81.7±0.35 | 82.9±0.18 | 81.9±0.26 | 76.5±0.45 |
| w/o tri-multi | 80.7±0.75 | 78.8±0.34 | 76.8±0.33 | 78.2±0.31 | 77.1±0.19 | 70.5±0.25 |
| w/o PPLM | 82.2±0.95 | 80.7±0.44 | 78.7±0.49 | 79.9±0.44 | 79.0±0.47 | 73.3±0.35 |
| w/o MSA | 81.3±0.38 | 79.2±0.76 | 76.5±0.38 | 78.3±0.31 | 76.9±0.35 | 70.3±0.22 |
| w/o Mdist | 67.4±0.95 | 66.7±0.93 | 63.0±0.91 | 65.6±0.81 | 63.5±1.00 | 53.9±0.51 |
| ***Heterodimer99 test set*** | | | | | | |
| PPLM-Contact | 63.6±1.14 | 59.9±1.87 | 55.9±1.04 | 59.4±2.37 | 58.0±1.47 | 49.7±1.02 |
| w/o cross-attn | 59.3±4.01 | 57.8±0.74 | 52.9±0.27 | 57.1±0.37 | 55.6±0.63 | 47.6±0.40 |
| w/o self-attn | 58.6±1.14 | 57.0±1.14 | 53.4±0.82 | 56.9±1.11 | 55.4±1.08 | 48.0±1.28 |
| w/o tri-multi | 51.5±1.98 | 50.7±3.18 | 46.5±1.62 | 50.4±2.27 | 48.8±1.99 | 40.3±1.30 |
| w/o PPLM | 54.9±2.88 | 54.4±1.49 | 50.5±0.76 | 54.3±1.57 | 52.8±1.23 | 45.3±0.50 |
| w/o MSA | 53.5±3.96 | 50.3±1.89 | 45.7±1.56 | 50.3±1.71 | 48.5±1.92 | 40.4±1.53 |
| w/o Mdist | 43.1±2.88 | 40.6±0.84 | 37.4±0.57 | 40.8±1.58 | 40.0±1.23 | 32.9±0.63 |

**Table S14.** Precision (%) of inter-protein contact prediction for PPLM-Contact2, AlphaFold2.3, AlphaFold3, and DMFold on all 343 homodimer and 119 heterodimer proteins. *L* denotes the length of the shorter protein in the dimer.

| Method | Top 1 | Top 10 | Top 50 | Top *L*/10 | Top *L*/5 | Top *L* |
|---|---|---|---|---|---|---|
| ***434 homodimers*** | | | | | | |
| PPLM-Contact2 | 90.7 | 90.4 | 89.0 | 90.1 | 87.9 | 85.1 |
| AlphaFold2.3 | 84.0 | 83.8 | 84.3 | 83.9 | 83.6 | 81.1 |
| AlphaFold3 | 79.9 | 81.6 | 81.5 | 81.0 | 80.6 | 78.3 |
| DMFold | 84.0 | 83.6 | 83.7 | 83.5 | 82.9 | 80.5 |
| ***119 heterodimers*** | | | | | | |
| PPLM-Contact2 | 94.1 | 92.4 | 91.1 | 91.9 | 90.8 | 88.0 |
| AlphaFold2.3 | 83.2 | 84.8 | 84.8 | 85.9 | 84.1 | 81.8 |
| AlphaFold3 | 84.9 | 86.1 | 86.2 | 86.6 | 85.7 | 83.4 |
| DMFold | 84.0 | 84.7 | 84.7 | 84.7 | 83.9 | 81.5 |

**Table S15.** Comprehensive statistical comparison between PPLM-Contact2 and complex structure-based models (AlphaFold2.3, AlphaFold3, and DMFold). For each dataset and baseline, improvements in per-target top *L* contact precision were evaluated using mean difference (Δ), relative percent gain, paired effect size (Cohen's d), Wilcoxon signed-rank *p*-values, and Benjamini–Hochberg FDR-adjusted *q*-values. Statistically significant results ($q < 0.05$) are indicated.

| Dataset | Mean Δ | Percent gain (%) | Effect size (Cohen's d) | Wilcoxon p-value | FDR q-value | Significant (q < 0.05) |
|---|---|---|---|---|---|---|
| ***vs AlphaFold2.3*** | | | | | | |
| Homodimers | 3.93 | 0.05 | 0.26 | 2.42E-18 | 4.84E-18 | ✓ |
| Heterodimers | 6.20 | 0.08 | 0.36 | 2.01E-10 | 2.01E-10 | ✓ |
| ***vs AlphaFold3*** | | | | | | |
| Homodimers | 6.77 | 0.09 | 0.34 | 1.62E-23 | 3.24E-23 | ✓ |
| Heterodimers | 4.65 | 0.06 | 0.25 | 4.60E-05 | 4.60E-05 | ✓ |
| ***vs DMFold*** | | | | | | |
| Homodimers | 4.53 | 0.06 | 0.28 | 1.34E-23 | 2.68E-23 | ✓ |
| Heterodimers | 6.57 | 0.08 | 0.40 | 5.51E-11 | 5.51E-11 | ✓ |

**Table S16.** Precision of interface residue identification by PPLM-Contact, DeepInter, CDPred, DeepHomo2.0 and GLINTER on the 343 homodimer and 119 heterodimer test proteins, using AlphaFold2-predicted monomer structures. The improvements of PPLM-Conatct over other beseline methods were evaluated using mean difference (Δ), relative percent gain, paired effect size (Cohen's d), Wilcoxon signed-rank *p*-values, and Benjamini–Hochberg FDR-adjusted *q*-values. Statistically significant results ($q < 0.05$) are indicated.

| Dataset | Average | Median | Mean Δ | Percent gain (%) | Effect size (Cohen's d) | Wilcoxon p-value | FDR q-value | Significant (q < 0.05) |
|---|---|---|---|---|---|---|---|---|
| ***Homodimer*** | | | | | | | | |
| PPLM-Contact | 0.824 | 0.916 | - | - | - | - | - | - |
| DeepInter | 0.784 | 0.892 | 0.039 | 5.0 | 0.253 | 7.82E-16 | 1.56E-15 | ✓ |
| PLMGraph-Inter | 0.765 | 0.847 | 0.058 | 7.6 | 0.408 | 1.75E-21 | 3.50E-21 | ✓ |
| CDPred | 0.782 | 0.869 | 0.041 | 5.3 | 0.173 | 3.29E-07 | 3.29E-07 | ✓ |
| DeepHomo2.0 | 0.691 | 0.779 | 0.133 | 19.3 | 0.622 | 3.51E-34 | 3.51E-34 | ✓ |
| GLINTER | 0.640 | 0.679 | 0.184 | 28.7 | 0.990 | 8.31E-43 | 1.66E-42 | ✓ |
| ***Heterodimer*** | | | | | | | | |
| PPLM-Contact | 0.695 | 0.752 | - | - | - | - | - | - |
| DeepInter | 0.620 | 0.649 | 0.075 | 12.1 | 0.370 | 2.04E-04 | 2.04E-04 | ✓ |
| PLMGraph-Inter | 0.649 | 0.674 | 0.046 | 7.1 | 0.288 | 2.12E-04 | 2.12E-04 | ✓ |
| CDPred | 0.608 | 0.679 | 0.087 | 14.2 | 0.409 | 1.63E-05 | 1.63E-05 | ✓ |
| GLINTER | 0.497 | 0.486 | 0.198 | 39.8 | 1.080 | 7.45E-18 | 7.45E-18 | ✓ |

**Table S17.** Precision of interface residue identification by PPLM-Contact, DeepInter, CDPred, DeepHomo2.0 and GLINTER on the 343 homodimer and 119 heterodimer test proteins, using experimental monomer structures. The improvements of PPLM-Conatct over other beseline methods were evaluated using mean difference (Δ), relative percent gain, paired effect size (Cohen's d), Wilcoxon signed-rank *p*-values, and Benjamini–Hochberg FDR-adjusted *q*-values. Statistically significant results ($q < 0.05$) are indicated.

| Dataset | Average | Median | Mean Δ | Percent gain (%) | Effect size (Cohen's d) | Wilcoxon p-value | FDR q-value | Significant (q < 0.05) |
|---|---|---|---|---|---|---|---|---|
| ***Homodimer*** | | | | | | | | |
| PPLM-Contact | 0.860 | 0.945 | - | - | - | - | - | - |
| PLMGraph-Inter | 0.810 | 0.918 | 0.051 | 6.3 | 0.331 | 1.54E-24 | 3.07E-24 | ✓ |
| DeepInter | 0.795 | 0.855 | 0.066 | 8.3 | 0.469 | 1.32E-28 | 2.64E-28 | ✓ |
| CDPred | 0.805 | 0.877 | 0.055 | 6.9 | 0.245 | 2.25E-13 | 4.50E-13 | ✓ |
| DeepHomo2.0 | 0.720 | 0.796 | 0.267 | 44.1 | 1.164 | 5.61E-19 | 5.61E-19 | ✓ |
| GLINTER | 0.651 | 0.696 | 0.210 | 32.2 | 1.197 | 1.75E-48 | 3.49E-48 | ✓ |
| ***Heterodimer*** | | | | | | | | |
| PPLM-Contact | 0.729 | 0.800 | - | - | - | - | - | - |
| PLMGraph-Inter | 0.633 | 0.679 | 0.097 | 0.153 | 0.458 | 2.06E-07 | 2.06E-07 | ✓ |
| DeepInter | 0.678 | 0.732 | 0.051 | 0.076 | 0.308 | 9.47E-05 | 9.47E-05 | ✓ |
| CDPred | 0.607 | 0.648 | 0.123 | 0.203 | 0.629 | 4.50E-10 | 4.50E-10 | ✓ |
| GLINTER | 0.498 | 0.515 | 0.231 | 0.464 | 1.224 | 6.40E-19 | 6.40E-19 | ✓ |

**Table S18.** Precision of interface residue identification by PPLM-Contact2, AlphaFold2.3, AlphaFold3, and DMFold. The improvements of PPLM-Conatct2 over other beseline methods were evaluated using mean difference (Δ), relative percent gain, paired effect size (Cohen's d), Wilcoxon signed-rank *p*-values, and Benjamini–Hochberg FDR-adjusted *q*-values. Statistically significant results ($q < 0.05$) are indicated.

| Dataset | Average | Median | Mean Δ | Percent gain (%) | Effect size (Cohen's d) | Wilcoxon p-value | FDR q-value | Significant (q < 0.05) |
|---|---|---|---|---|---|---|---|---|
| ***Homodimer*** | | | | | | | | |
| PPLM-Contact2 | 0.897 | 0.962 | - | - | - | - | - | - |
| AlphaFold2.3 | 0.888 | 0.957 | 0.008 | 1.0 | 0.082 | 1.15E-04 | 1.15E-04 | ✓ |
| AlphaFold3 | 0.873 | 0.953 | 0.024 | 2.8 | 0.180 | 1.42E-07 | 2.83E-07 | ✓ |
| DMFold | 0.879 | 0.955 | 0.018 | 2.0 | 0.181 | 4.93E-10 | 9.85E-10 | ✓ |
| ***Heterodimer*** | | | | | | | | |
| PPLM-Contact2 | 0.904 | 0.945 | - | - | - | - | - | - |
| AlphaFold2.3 | 0.874 | 0.930 | 0.030 | 3.5 | 0.328 | 2.45E-07 | 4.90E-07 | ✓ |
| AlphaFold3 | 0.887 | 0.941 | 0.017 | 1.9 | 0.162 | 2.04E-02 | 2.04E-02 | ✓ |
| DMFold | 0.879 | 0.930 | 0.025 | 2.8 | 0.317 | 2.12E-07 | 2.12E-07 | ✓ |

**Table S19.** Detailed information on the PPI datasets. The numbers in parentheses represent the original count of negative samples, while the numbers outside the parentheses indicate the actual count of negative samples after removing anomalous samples.

| Dataset | Name | Species | Number of positive samples | Number of negative samples |
|---|---|---|---|---|
| Training set | Human | *H. sapiens* | 43,138 | 430,947 (431,379) |
| Test sets | Mouse | *M. musculus* | 5,000 | 49,999 (50,000) |
| | Fly | *D. melanogaster* | 5,000 | 49,993 (50,000) |
| | Worm | *C. elegans* | 5,000 | 49,996 (50,000) |
| | Yeast | *S. cerevisiae* | 5,000 | 49,963 (50,000) |
| | Ecoli | *E. coli* | 2,000 | 16,232 (20,000) |

**Table S20.** Dimensional transformations during feature extraction and processing in the PPLM-PPI. $L_1$ and $L_2$ present the lengths of the two proteins, respectively.

| Feature Type | Initial Dimension | After Pooling (Max/Mean) | After Linear layer |
|---|---|---|---|
| **Embedding** | ($L_1$×1280), ($L_2$×1280) | 1280, 1280 | 660, 660 |
| **Intra-attention** | ($L_1$×$L_1$×660), ($L_2$×$L_2$×660) | 660, 660 | 660, 660 |
| **Inter-attention** | ($L_1$×$L_2$×660) | 660 | 660 |

**Table S21.** Dimensional transformations through the multi-layer perceptron (MLP) in the PPLM-PPI.

| Layer | Input Dimension | Output Dimension | Activation Function | Normalization |
|---|---|---|---|---|
| **Linear 1** | 660×10 | 1024 | ReLU | LayerNorm |
| **Linear 2** | 1024 | 512 | ReLU | LayerNorm |
| **Linear 3** | 512 | 256 | ReLU | LayerNorm |
| **Linear 4** | 256 | 128 | ReLU | LayerNorm |
| **Linear 5** | 128 | 1 | Sigmoid | LayerNorm |

**Table S22.** Description of features used in PPLM-Contact. $L_1$ and $L_2$ present the lengths of the two proteins, respectively.

| Type | Feature | Dimension |
|---|---|---|
| PPLM | Inter-protein attention | ($L_1$, $L_2$, 660) |
| MSA | Monomer PSSM | ($L_1$, 20), ($L_2$, 20) |
| | Monomer DCA | ($L_1$, $L_1$, 2), ($L_2$, $L_2$, 2) |
| | Monomer MSA embedding | ($L_1$, 768), ($L_2$, 768) |
| | Monomer MSA attention | ($L_1$, $L_1$, 144), ($L_2$, $L_2$, 144) |
| | Inter-protein DCA | ($L_1$, $L_2$, 2) |
| | Inter-protein MSA attention | ($L_1$, $L_2$, 144) |
| Monomer structure | Monomer distance map | ($L_1$, $L_1$, 64), ($L_2$, $L_2$, 64) |

### Supplementary Figures

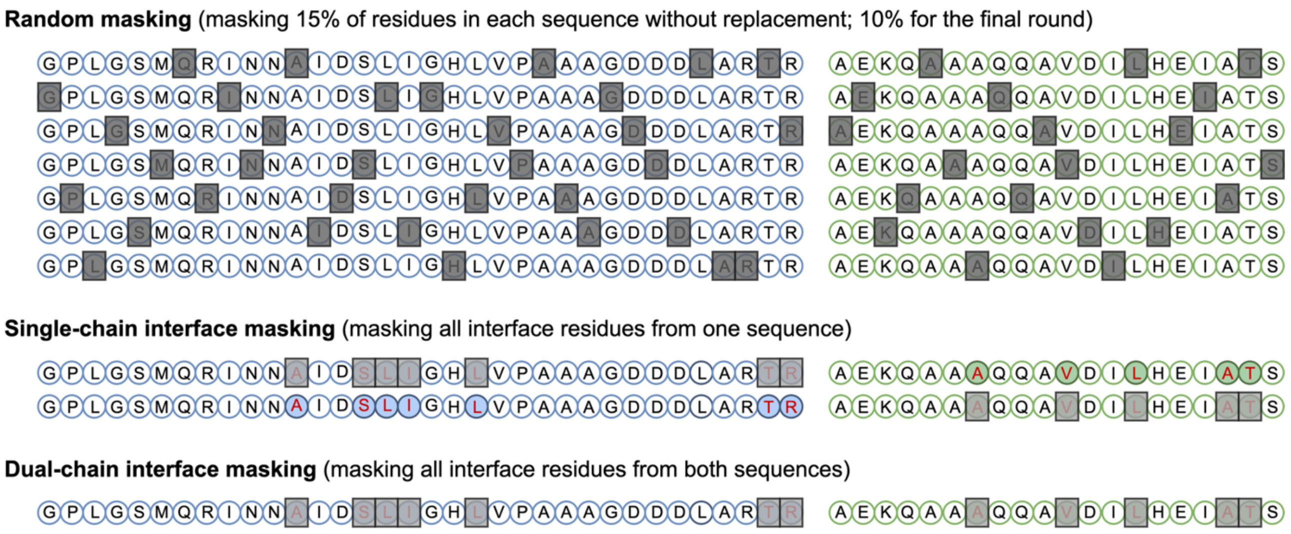


**Figure S1. Schematic of masking strategies for perplexity calculation.** We designed three masking strategies. (i) Random masking: randomly masking 15% of residues in each sequence without replacement (10% for the final round), with the average perplexity across rounds used as the perplexity for the sequence pair. (ii) Single-chain interface masking: masking all interface residues from one sequence at a time. (iii) Dual-chain interface masking: masking all interface residues from both sequences simultaneously. Interface residues are highlighted using red text and colored backgrounds.

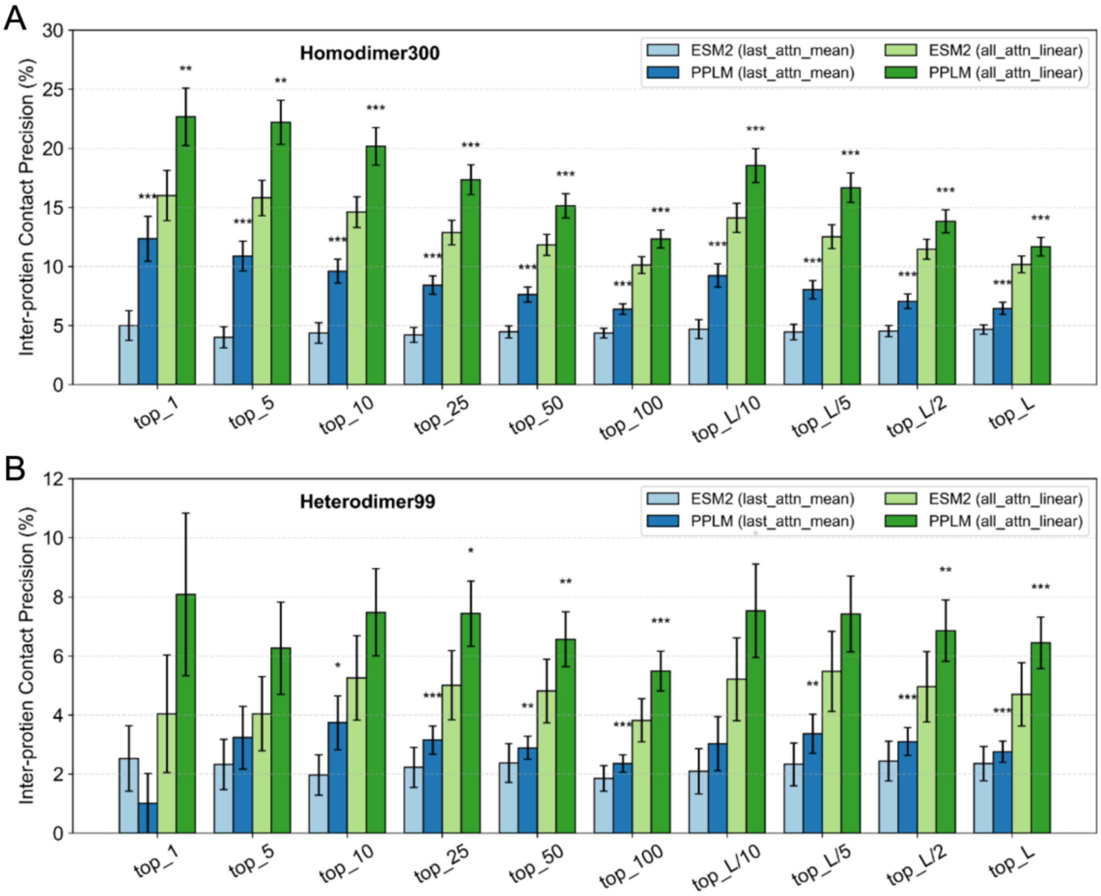


**Figure S2. Comparative analysis of inter-protein attentions from PPLM and ESM2.** (A) Results on the Homodimer300 test set. (B) Results on the Heterodimer99 test set. Bar plots show Top-k inter-protein contact precision (%) for PPLM and ESM2 under two settings: unsupervised (last-layer mean attention) and linear-probe (all-layer attention + frozen linear layer). Error bars indicate standard errors (SE) across proteins. Asterisks denote significance of paired Wilcoxon tests (one-sided, PPLM > ESM2): $p < 0.05$ (*), p < 0.01 (**), and $p < 0.001$ (***).

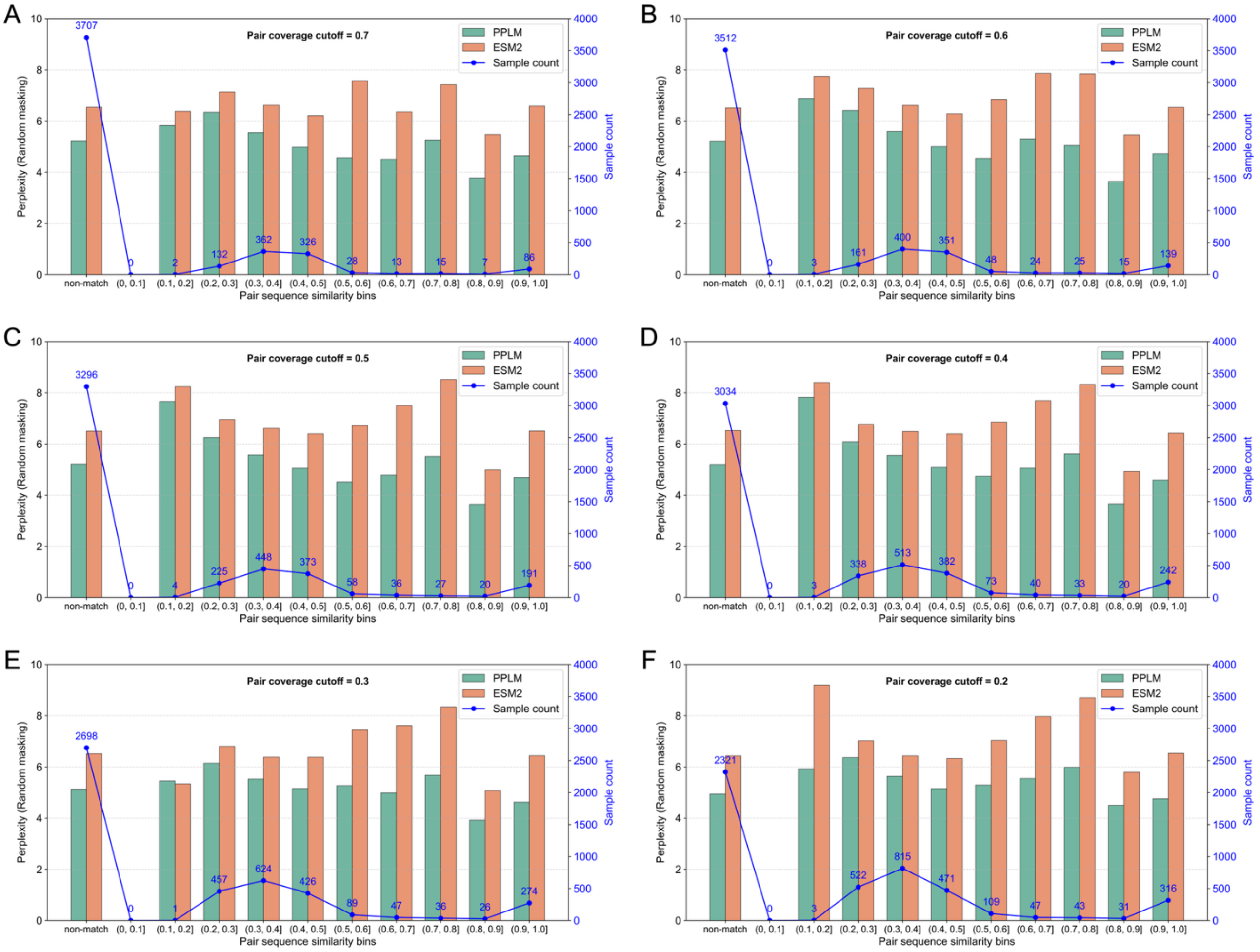


**Figure S3. Sequence pair identity distribution and model performance across coverage cutoffs.** Distribution of pairwise sequence identities between validation and training pairs, and corresponding model performance (perplexity) under alignment coverage cutoffs of 0.7 (**A**), 0.6 (**B**), 0.5 (**C**), 0.4 (**D**), 0.3 (**E**), and 0.2 (**F**).

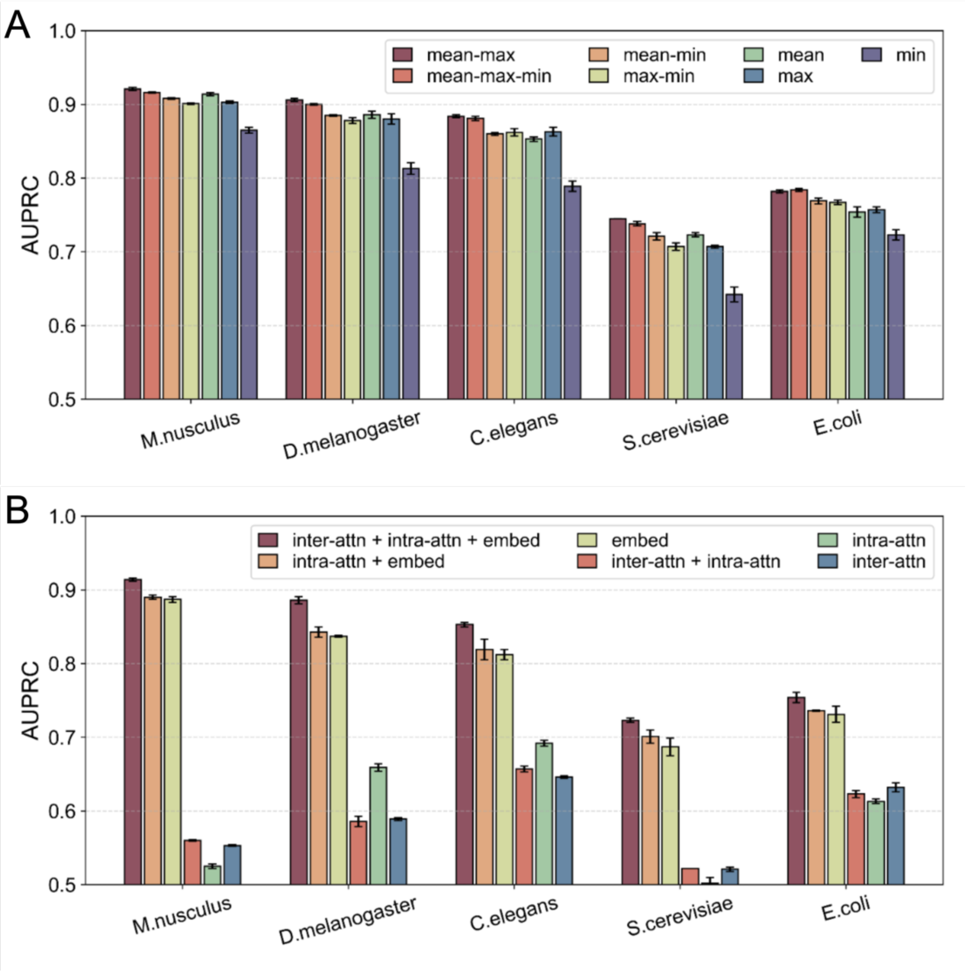


**Figure S4. Performance of PPLM-PPI and its Ablated variants. (A)** AUPRC of different pooling strategies **(B)** AUPRC of different feature combinations. Values represent the mean ± 95 % confidence intervals (CI) computed across three random seeds.

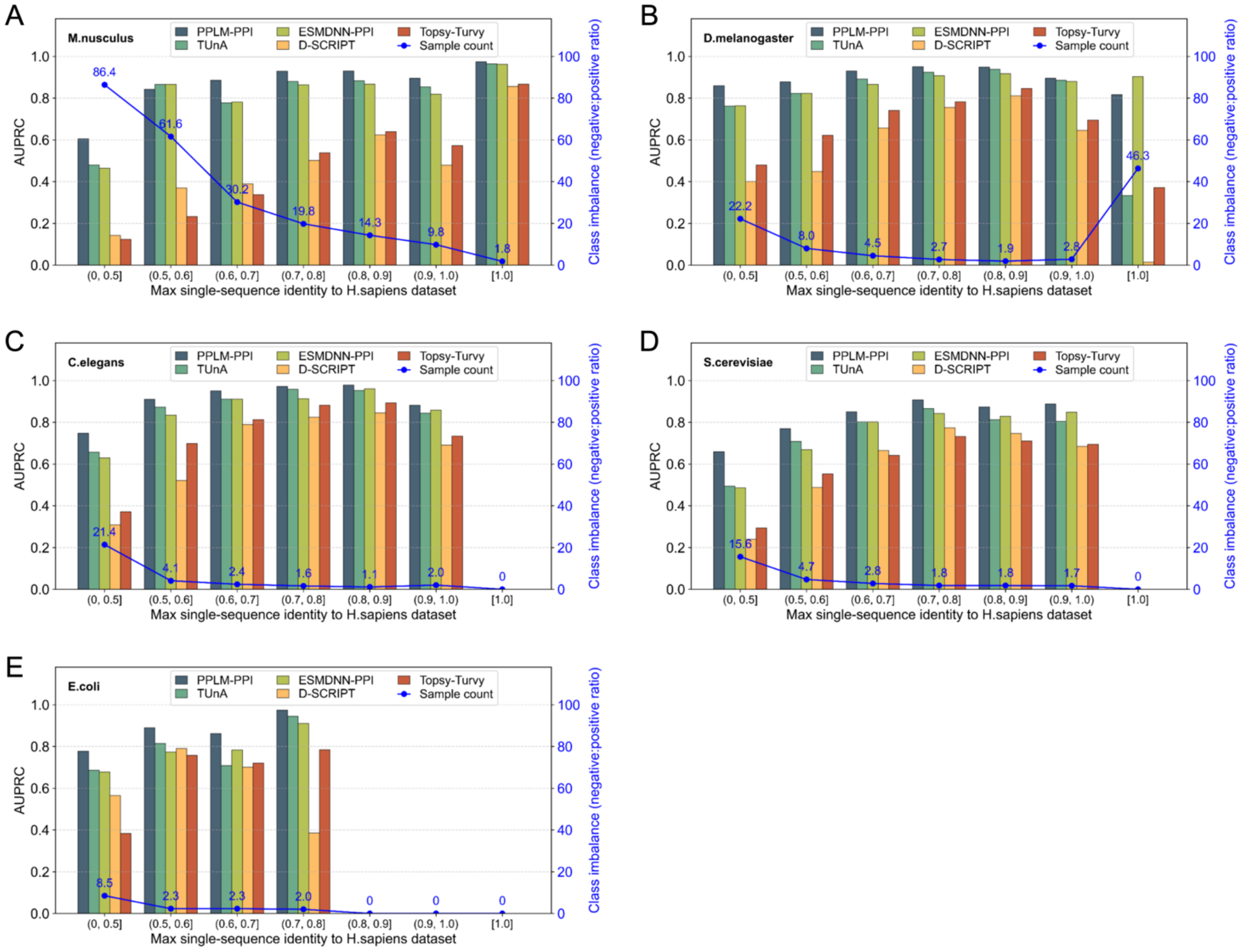


**Figure S5. AUPRC of PPLM-PPI and baseline methods across sequence-identity bins for five species test sets. (A)** *M. musculus*. **(B)** *D. melanogaster*. **(C)** *C. elegans*. **(D)** *S. cerevisiae*. **(E)** *E. coli*.

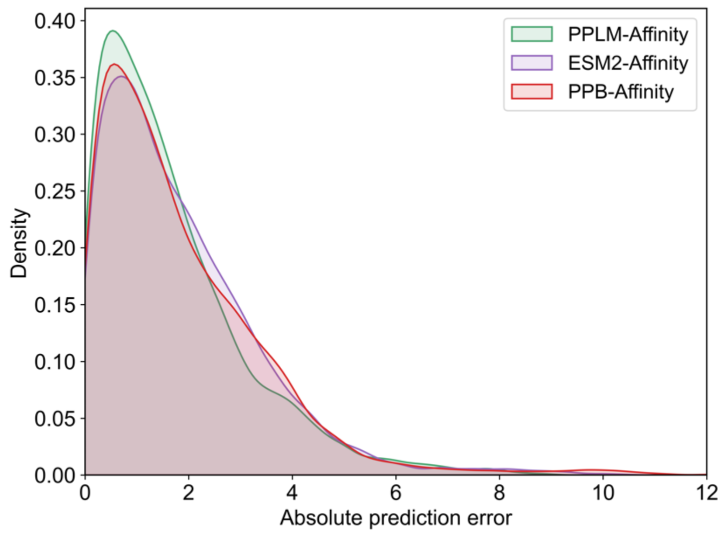


**Figure S6. Distribution of absolute errors between predicted and experimental binding affinities.**

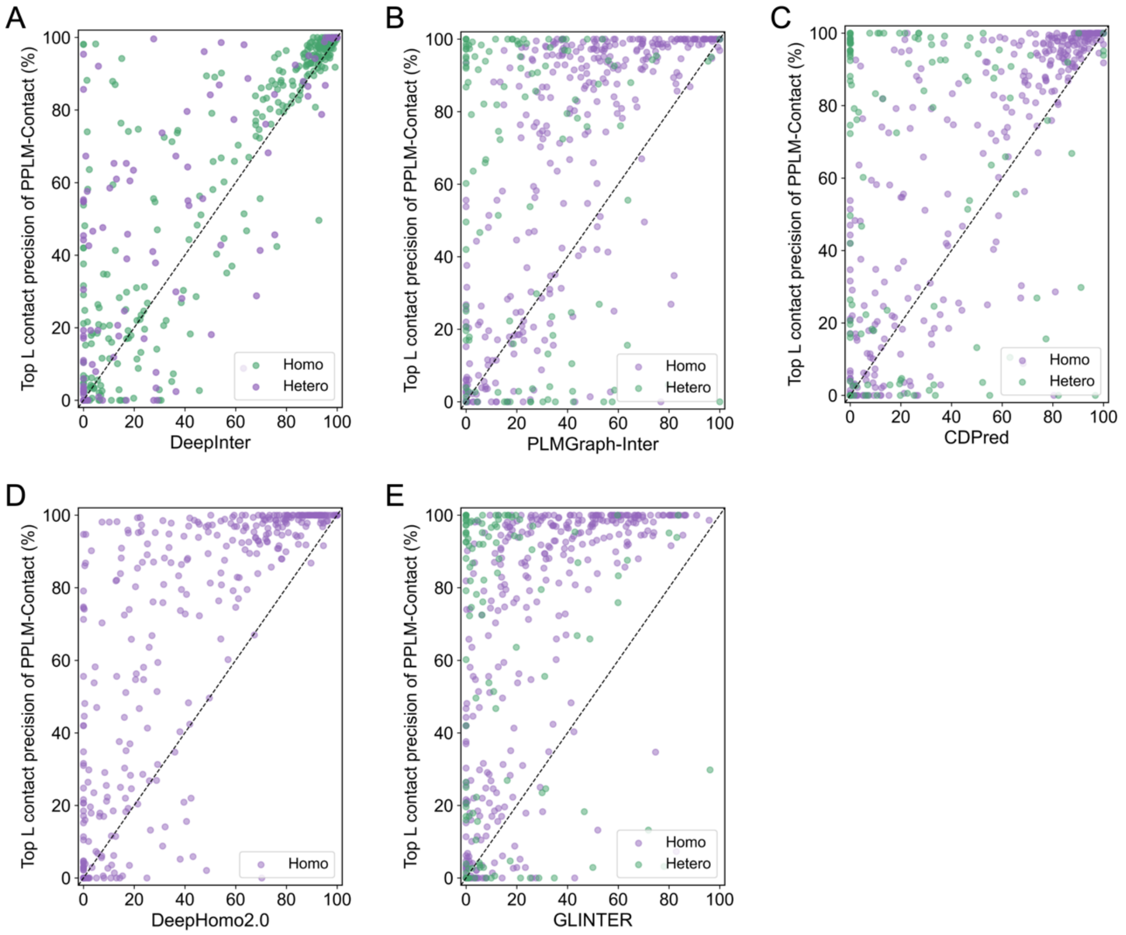


**Figure S7. Head-to-head comparison of top *L* contact precision of PPLM-Contact to each baseline methods using AlphaFold2-predicted monomer structures. (A)** vs DeepInter. **(B)** vs PLMGraph-Inter. **(C)** vs CDPred. **(D) vs DeepHomo2.0. (E)** vs GLINTER.

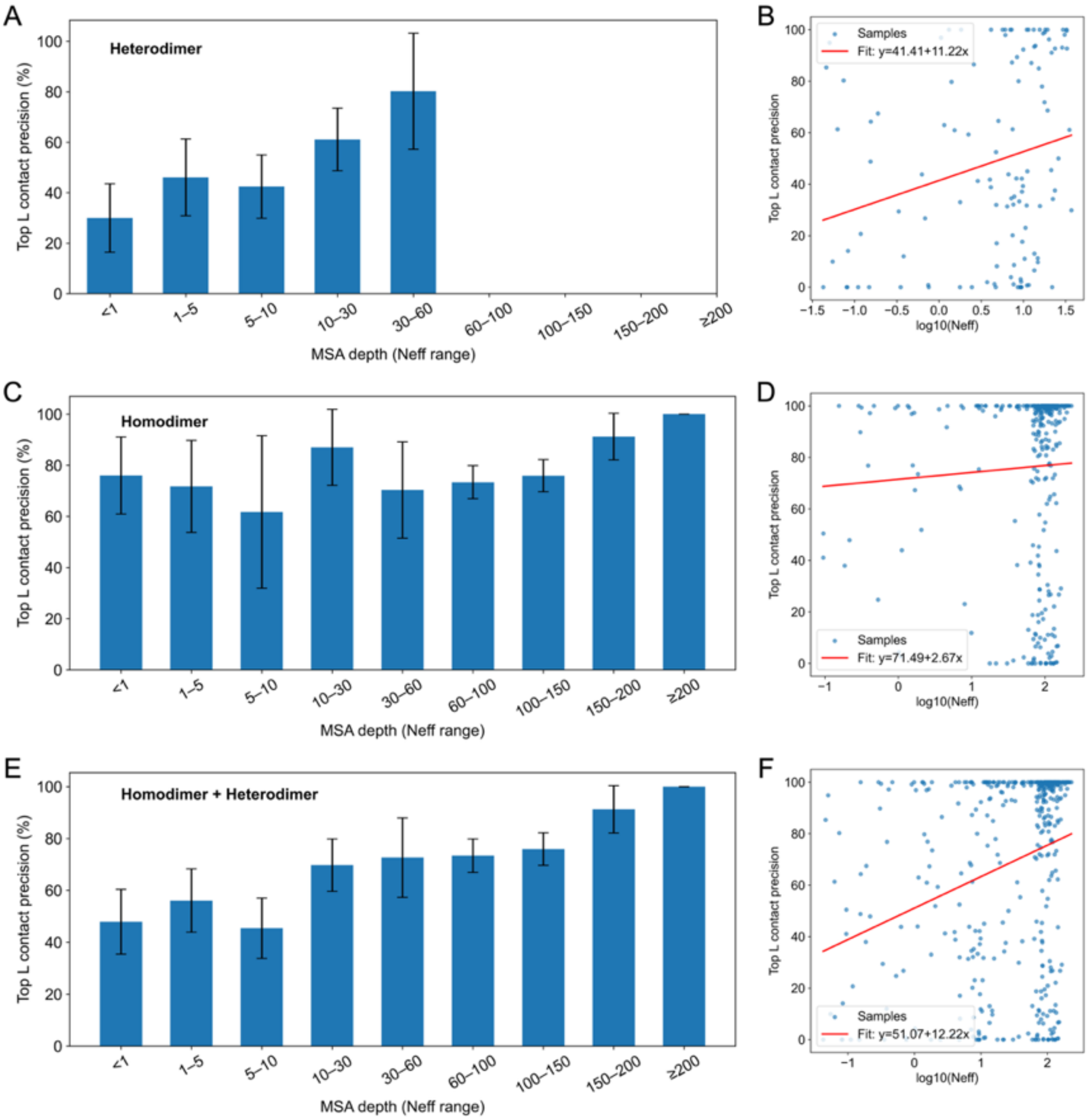


**Figure S8. Relationship between MSA depth and inter-protein contact-prediction precision for PPLM-Contact.** (A–B) Heterodimers, (C–D) homodimers, and (E–F) all test complexes. Bars show mean top-L precision across MSA-depth bins with 95% confidence intervals, and scatter plots display precision versus $\log_{10}$(*Neff*).

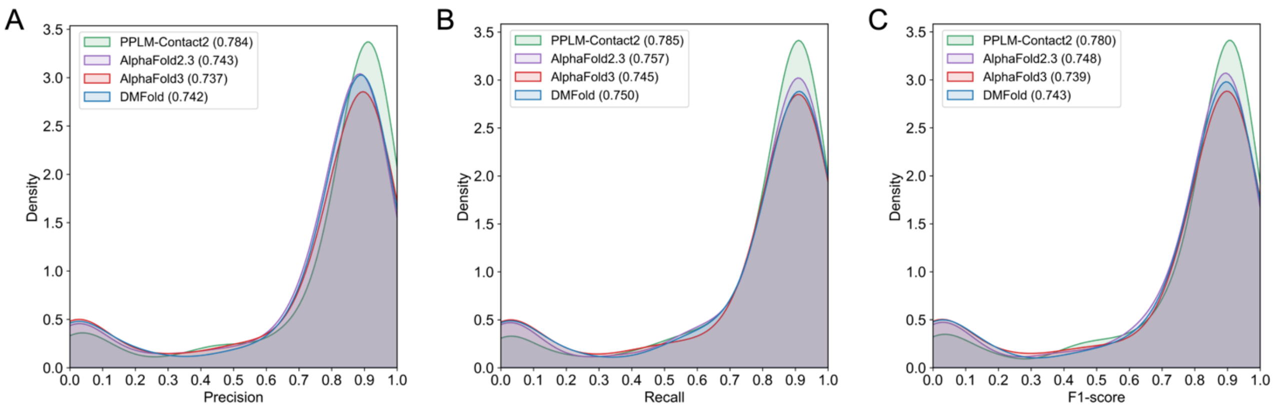


**Figure S9. Distribution of precision (A), recall (B), and F1-score (C) for inter-protein contact prediction by PPLM-Contact2, AlphaFold2.3, AlphaFold3, and DMFold on all 462 test proteins.** A contact probability threshold of 0.4 is used for PPLM-Contact2. For control methods, inter-protein contacts are extracted from their predicted complex structures. Values indicate the average precision, recall, and F1-score for each method.

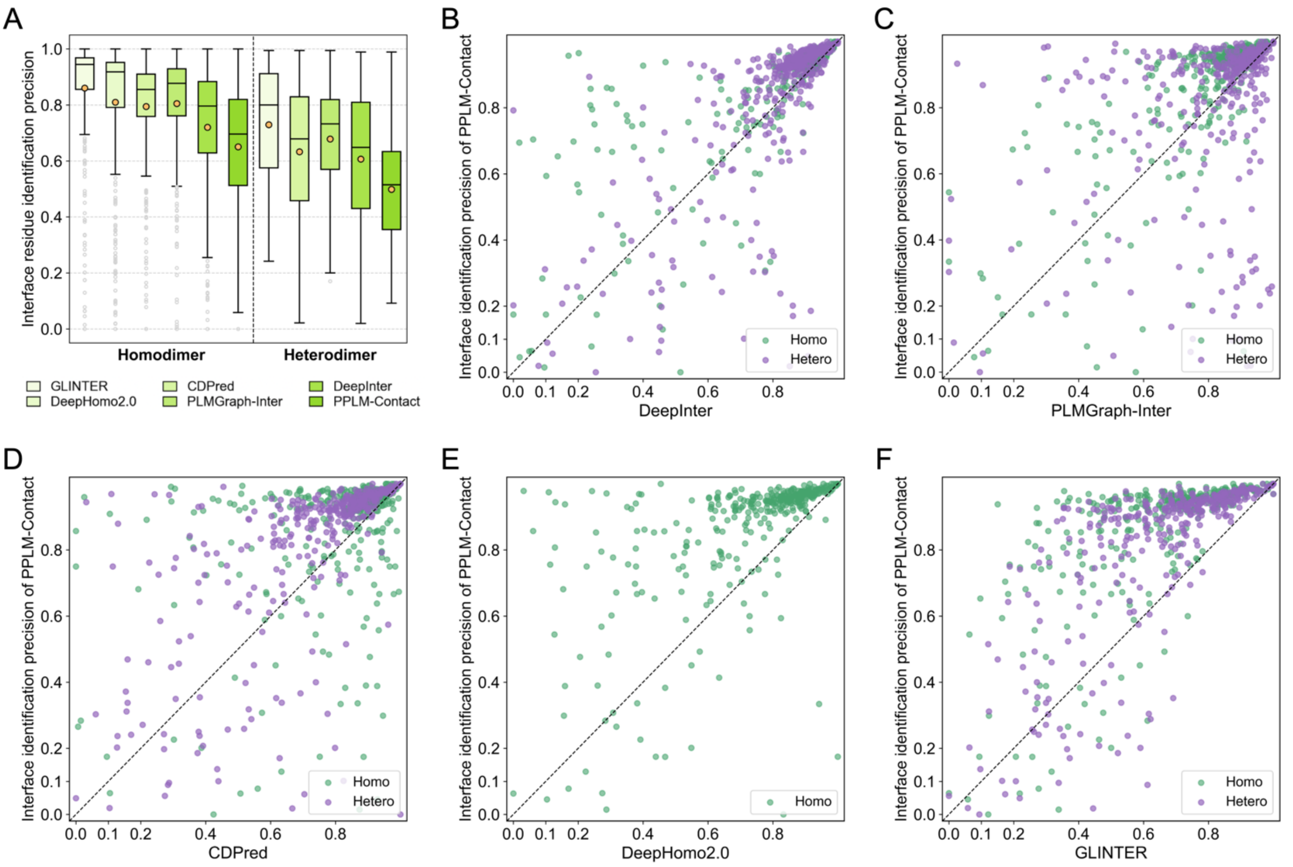


**Figure S10. Comparative analysis of protein–protein interface residue identification using experimental monomer structures. (A)** Precision of interface-residue identification by general contact-prediction methods (GLINTER, DeepHomo2.0, PLMGraph-Inter, CDPred, DeepInter, and PPLM-Contact). **(B–F)** Head-to-head comparison of identification precision between PPLM-Contact and each control method on the homodimer and heterodimer proteins, respectively.

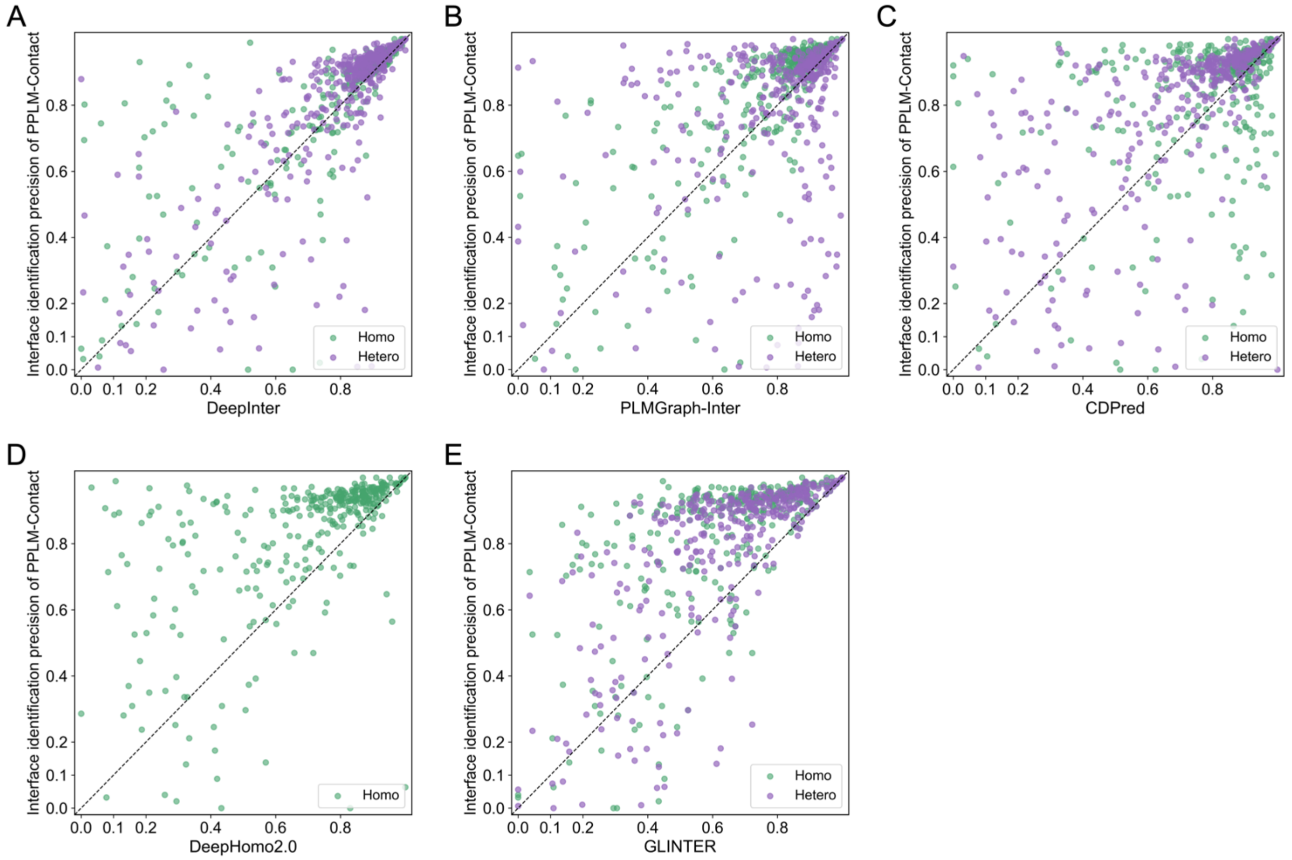


**Figure S11. Head-to-head comparison of interface residue identification precision between PPLM-Contact and each baseline methods using AlphaFold2-predicted monomer structures for interface residue identification. (A)** vs DeepInter. **(B)** vs PLMGraph-Inter. **(C)** vs CDPred. **(D)** vs DeepHomo2.0. **(E)** vs GLINTER.

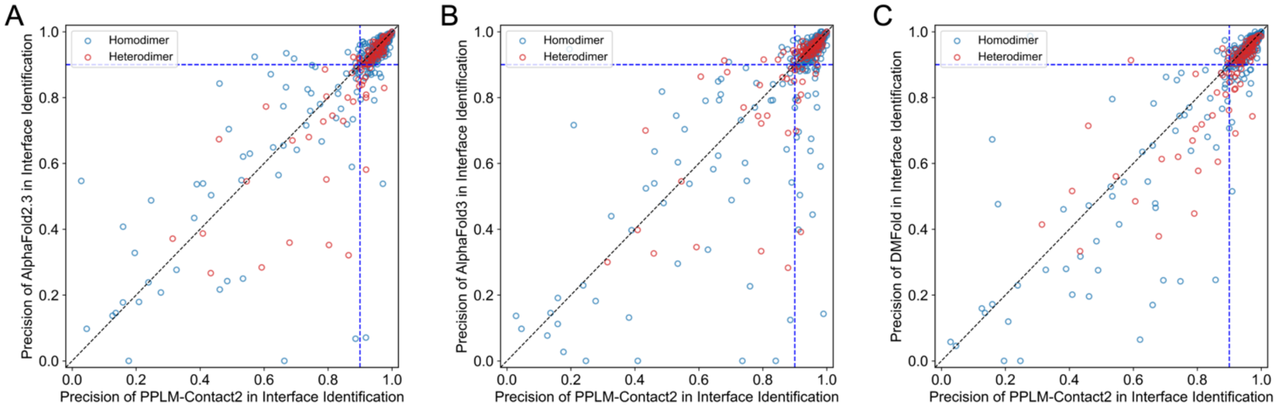


**Figure S12. Head-to-head comparison of interface residue identification precision between PPLM-Contact2 and complex structure-based methods. (A)** vs AlphaFold2.3. **(B)** vs AlphaFold3**. (C)** vs DMFold**.**

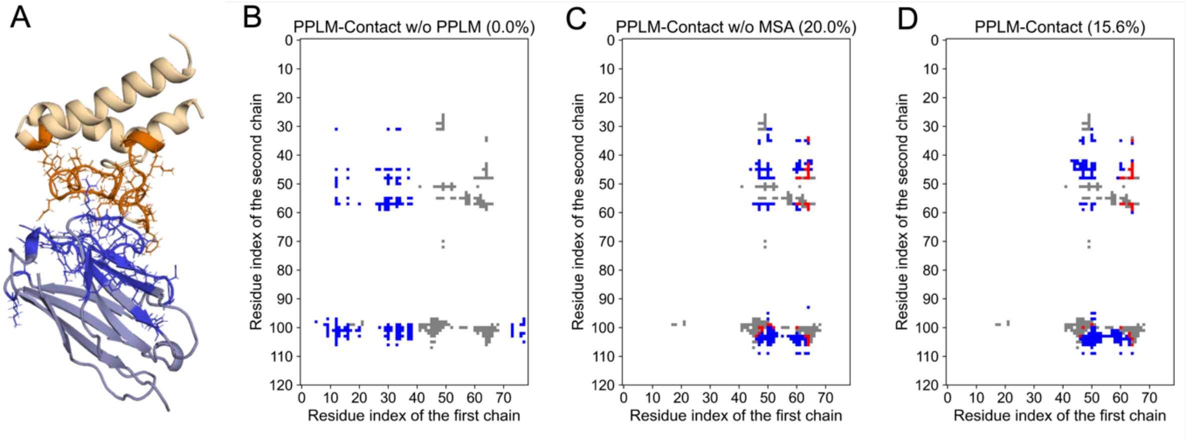


**Figure S13. A representative challenging heterodimer for inter-protein contact prediction: the EC2 domain of CD9 in complex with nanobody 4C8 (PDB ID: 6Z20). (A)** Structural visualization of the complex with interface residues highlighted. **(B-C)** Inter-protein contact maps predicted by PPLM-Contact w/o PPLM, PPLM-Contact w/o MSA, and the full PPLM-Contact model, respectively. In each panel, gray dots indicate ground-truth contacts, red dots denote correctly predicted contacts (true positives), and blue dots represent false positives. For each method, the number in parentheses indicates the top $N$ contact precision, where $N$ equals the number of ground-truth contacts.

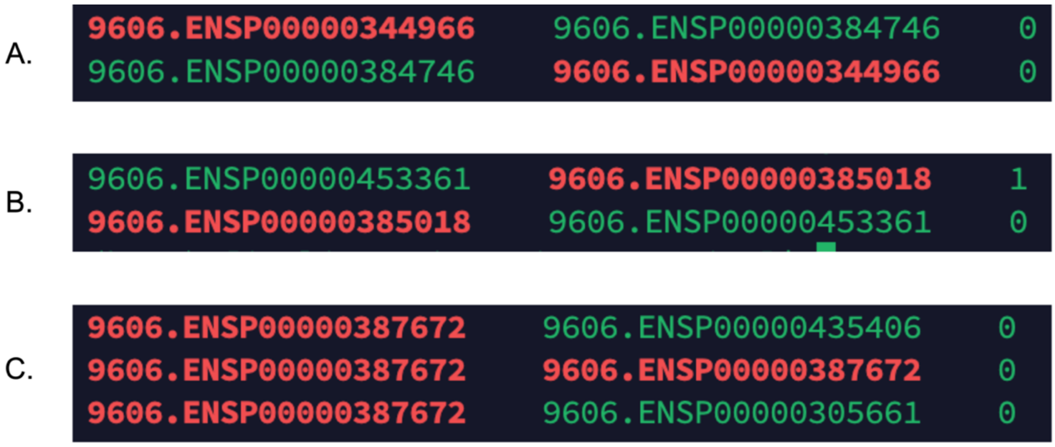


**Figure S14. Example of anomalous samples observed in *H. sapiens* dataset.** (A) Duplicate, (B) Erroneous, and (C) Invalid. Duplicate data: only one sample was retained. Erroneous data: only the positive sample was retained. All invalid samples were removed.

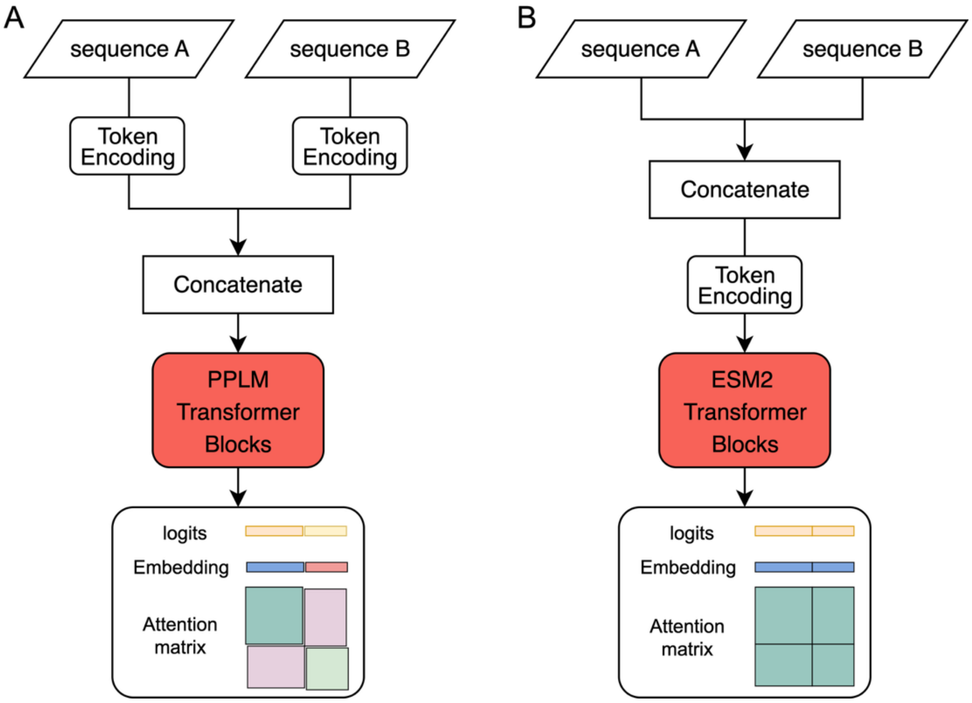


**Figure S15. Overview of PPLM and ESM2 processing for protein sequence pair.** (A) In PPLM, the two input sequences are first encoded separately and then concatenated before being passed through the model's pairwise transformer blocks. (B) In ESM2, which is designed for single-chain modeling, the two sequences are directly concatenated into a single input and processed by its single-chain transformer blocks. For both PPLM and ESM2, the outputs include logits, embdding and attention matrix corresponding to the input sequence pair.

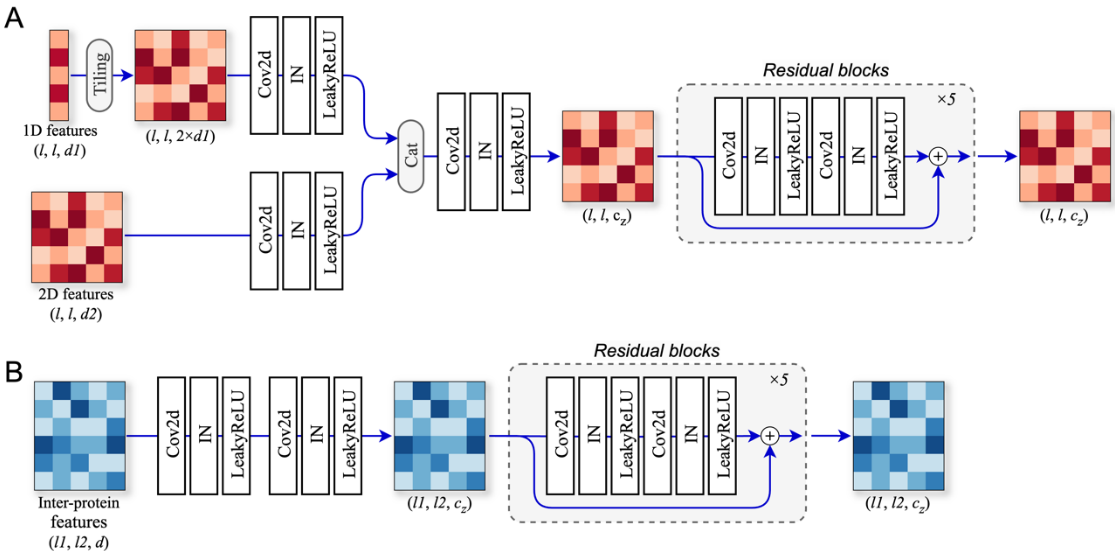


**Figure S16. Architectures of intra-protein ResNet (A) and inter-protein ResNet (B).** Here, *l*, *l1*, and *l2* denote the lengths of monomer proteins; *d1* and *d2* the dimensions of 1D and 2D features of monomer proteins, respectively; *d* the dimension of 2D features of inter-protein features; and $c_z$ the dimension of pair representation.

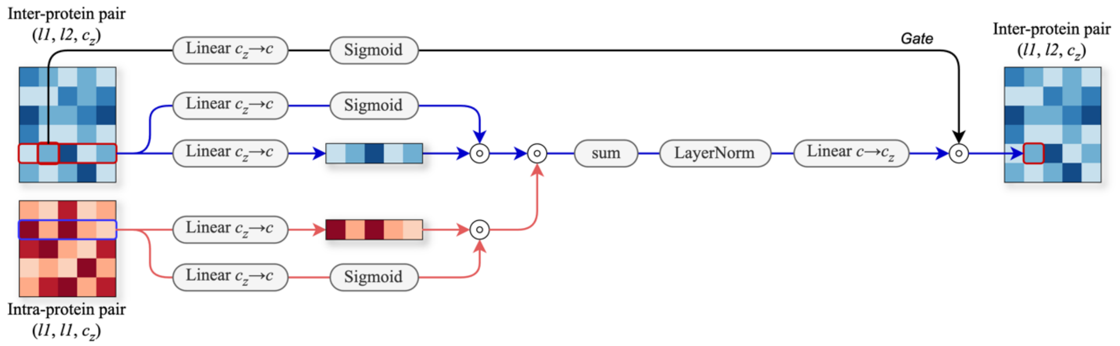


**Figure S17. Architecture of triangle multiplication module.** Here, *l1* and *l2* present lengths of monomer proteins; $c_z$ denotes the dimensions; and *c* indicates the number of channels.

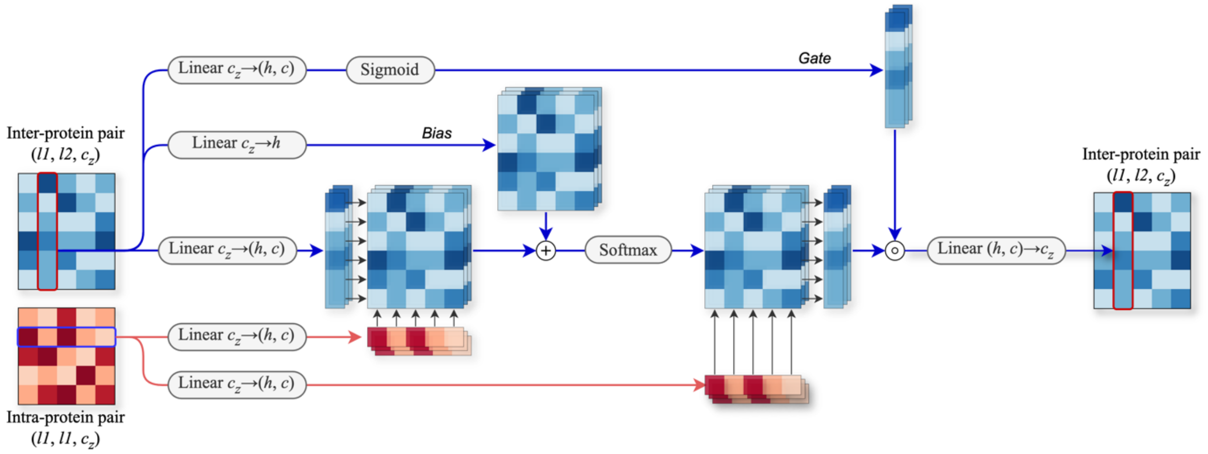


**Figure S18. Architecture of cross-attention module.** Here, *l1* and *l2* present lengths of monomer proteins; $c_z$ denotes the dimensions; *h* represents the number of attention heads; and *c* indicates the number of channels in each attention head.

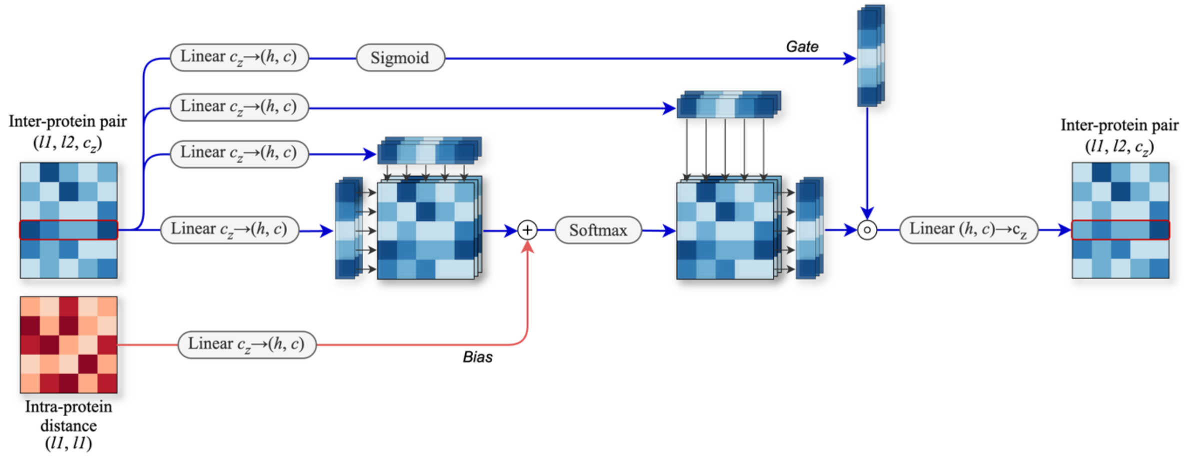


**Figure S19. Architecture of self attention module.** Here, *l1* and *l2* present lengths of monomer proteins; $c_z$ denotes the dimensions; *h* represents the number of attention heads; and *c* indicates the number of channels in each attention head.
